# A yeast mating-based platform enables the generation and screening of ultra large antibody libraries

**DOI:** 10.1101/2025.10.06.680701

**Authors:** Lester Frei, Sai T. Reddy

## Abstract

Yeast display is a powerful platform for antibody engineering, offering eukaryotic protein folding and compatibility with fluorescence-activated sorting (FACS). However, conventional yeast display faces limitations in terms of library size and display heterogeneity due to plasmid-based expression. Here, we present a yeast display platform that combines genomic library integration and high-efficiency mating to generate ultra-diverse antibody libraries approaching the diversity of phage-libraries (>10^11^). We engineered two yeast strains, LFYa and LFYalpha, featuring genomically integrated landing pads for high-efficiency library insertion. To enable compatibility with deep sequencing, we integrated the recombinase BxB1, which links heavy (HC) and light chain (LC) information onto a single chromosome. Using our platform, we constructed synthetic antibody libraries with HC and LC diversities of 4.1×10^7^ and 1.7×10^7^ variants, respectively and with an improved mating protocol, we achieved a combinatorial library diversity exceeding 10^11^. We demonstrated that genomic integration yields uniform surface display. Screening this antibody library against multiple antigens resulted in the discovery of binders with affinities in the single-digit nanomolar to picomolar range, demonstrating the platform’s utility for the discovery of antibodies with therapeutically relevant affinities. This work establishes a robust and deep sequencing-compatible yeast display system that overcomes key limitations of previous mating-based platforms.

## Introduction

Display platforms are powerful tools for directed evolution and protein engineering, enabling the high-throughput screening of libraries to isolate variants with enhanced properties, such as improved binding affinity or stability (1, 2). Due to their clinical importance, antibodies have been a primary focus, with display technologies facilitating the discovery of several approved therapeutics, including blockbusters such as adalimumab and dupilumab (3). While phage display remains widely used, *Saccharomyces cerevisiae*-based yeast surface display (YSD) offers distinct advantages for the engineering of complex proteins. Yeast combines the rapid growth rate and ease of manipulation typical of prokaryotic systems with the eukaryotic protein folding machinery, disulfide bond formation, and post-translational modifications critical for antibody functionality (4). These features make yeast display particularly well-suited for screening antibodies (fragment antigen-binding (Fab) or single-chain variable fragment (scFv) formats), as demonstrated by its success in optimizing affinity, stability, and expression yields of therapeutic candidates (4, 5).

However, a persistent limitation of yeast display is its smaller library size, typically constrained to 10^7^ - 10^9^ variants, several orders of magnitude smaller than phage or ribosome display libraries (10^11^ - 10^12^ and 10^12^ - 10^15^ respectively) (2). This bottleneck arises primarily from low transformation efficiencies in yeast as well as instability of plasmid-based libraries during propagation (6). To overcome this, yeast mating, a natural process wherein haploid MATa and MATα cells fuse to form MATa/α diploids has been employed to combinatorially shuffle antibody HC and LC libraries encoded in separate strains, which can result in library sizes exceeding 10^11^, rivaling phage display diversity (7, 8). During yeast mating, MATa and MATα cells secrete peptide pheromones (a- and α-factor). These pheromones bind to the G protein-coupled receptors (GPCRs) Ste2 and Ste3, on MATa and MATα cells respectively, initiating a signaling cascade that leads to G1 cell cycle arrest (9). The cells subsequently undergo shmooing, a process in which cells protrude towards each other until the high-affinity interaction between Aga2 (on MATa) and Sag1 (on MATα) physically connects them. This facilitates cell fusion and karyogamy which gives rise to a MATa/α diploid cell.(9, 10)

Despite its potential to generate large, shuffled libraries, existing mating-based approaches suffer from critical limitations: (i) reliance on single-copy number plasmids with the CEN/ARS origin of replication (ori) leads to rapid library loss due to uneven plasmid segregation; (ii) reported mating efficiencies remain suboptimal for generating highly diverse libraries; and (iii) the lack of a mechanism to physically link HC and LC sequences limits deep sequencing-based workflows (8). These issues have hindered the widespread adoption of yeast mating for large-scale antibody library generation and screening.

Here, we present a yeast display platform that addresses these challenges by integrating three key innovations. First, we engineered *S. cerevisiae* strains (LFYa and LFYalpha) with genomically encoded landing pads for stable, high-efficiency library integration, eliminating plasmid-associated instability. Second, we systematically optimized mating conditions to achieve significantly improved mating efficiency when compared to prior methods. Third, we incorporated the serine recombinase BxB1 to combine HC and LC information onto a single chromosome, enabling compatibility with deep sequencing workflows. Using this system, we constructed an antibody Fab library with an ultra large diversity (>10^11^), which following screening led to the discovery of binders against several therapeutic target antigens (TNFα, HER2, and Influenza HA). By decoupling library size from transformation efficiency and ensuring genomic stability, this platform expands the scope of yeast display for high-throughput protein engineering.

## Results

### Design and construction of yeast mating strains

As a basis for the platform, we utilized two yeast strains. The first strain, BJ5465 is a MATa strain and the parent strain for EBY100, which is commonly used for yeast display of antibody libraries because of its protease deficiency (11). The second strain, BJ5464 is a closely related MATα strain. In both strains, genomic deletions and integrations were introduced using the markerless yeast localization and overexpression (MyLO) CRISPR-Cas9 toolkit (**Table 1**, Methods) (12). Although they are naturally inactive in BJ5464 and BJ5465, *ura3* and *trp1* genes were completely removed in both strains so that they could be used as selection markers for antibody library integration while avoiding potential for spontaneous reversion of the inactive gene to an active form. Furthermore, *sag1* in BJ5465 was deleted by integrating *aga1* under the control of the Gal1 promoter (P_Gal1_). Expression of *sag1* is silenced in MATa cells and the integration of *aga1* under the control of P_Gal1_ is crucial for galactose inducible yeast display (13). Next, we integrated the gene for the well-tempered controller (WTC) into the FF18 locus of BJ5464. WTC is an anhydrotetracycline (aTc) inducible system and has been noted for its low basal expression level and for being strongly inducible (14), which we utilized to control the expression of the large serine recombinase BxB1.

**Table 1.**
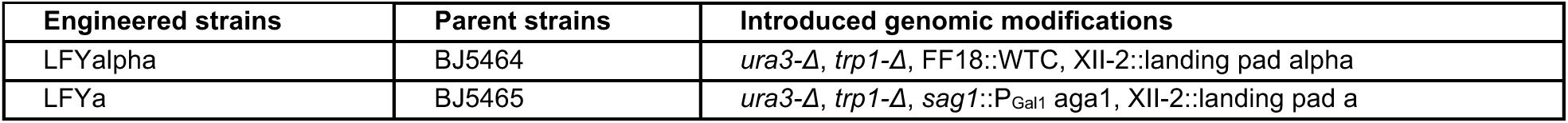
Genomic modifications introduced into BJ5464 and BJ5465. Sequence maps can be found in **Supplementary Material**.

In both strains, a landing pad was integrated into the XII-2 locus on chromosome (Chr) XII using the MyLO approach, and the resulting strains are referred to as LFYa and LFYalpha. The landing pads consist of the MFα1 terminator (T_MFα1_) and the open reading frame (ORF) of either *ura3* or *trp1*, which are inactive due to the lack of a promoter. An 18 basepair (bp) recognition site for the homing endonuclease I-SceI was placed between T_MFα1_ and *ura3*/*trp1* (**Fig. 1a, Supplementary Material**) (15). I-SceI is a yeast meganuclease that has been noted for its high specificity and activity in a range of different organisms (16, 17). For genomic integration, the payload is co-transformed with a linearized plasmid carrying the strong constitutive yeast promoter P_TDH3_, the ORF for I-SceI homing endonuclease and the strong yeast terminator T_TDH1_ (18). This plasmid is referred to as pI-SceI (**Fig. 1b**). Upon transformation, I-SceI is transiently expressed, leading to a double-stranded DNA (dsDNA) break in the landing pad, thus enabling efficient genomic integration via homology directed repair (HDR). The HDR template contains the promoter for *ura3/trp1*, P_Gal1_, HC/LC and is flanked by 60 bp homology to the ORF of *ura3*/*trp1* and T_MFα1_ (**Fig. 1b**). Successful genomic integration restores functionality of *ura3* and *trp1* in LFYa and LFYalpha respectively. The *ura3* and *trp1* markers are used to select and enrich cells that successfully genomically integrated the fragments (**Fig. 1c**). This strategy of only integrating the promoter for the selectable markers yielded better results than introducing the complete ORF. The latter approach leads to a substantial degree of background growth. In a subsequent step, the cells are mated to obtain diploid cells. The resulting diploid cells are double positive for *ura3* and *trp1* and carry an antibody HC and LC for surface display (**Fig. 1d**). When using different ratios of pI-SceI and HDR template, varying levels of integration efficiency were observed, with 1 pmol pI-SceI and 2 pmol of HDR template yielding approximately 7.5×10^6^ transformation events per cuvette (**Fig. 1e, Supplementary Fig. 1**). This transformation efficiency rivals plasmid-based library transformation protocols.

**Fig. 1.**
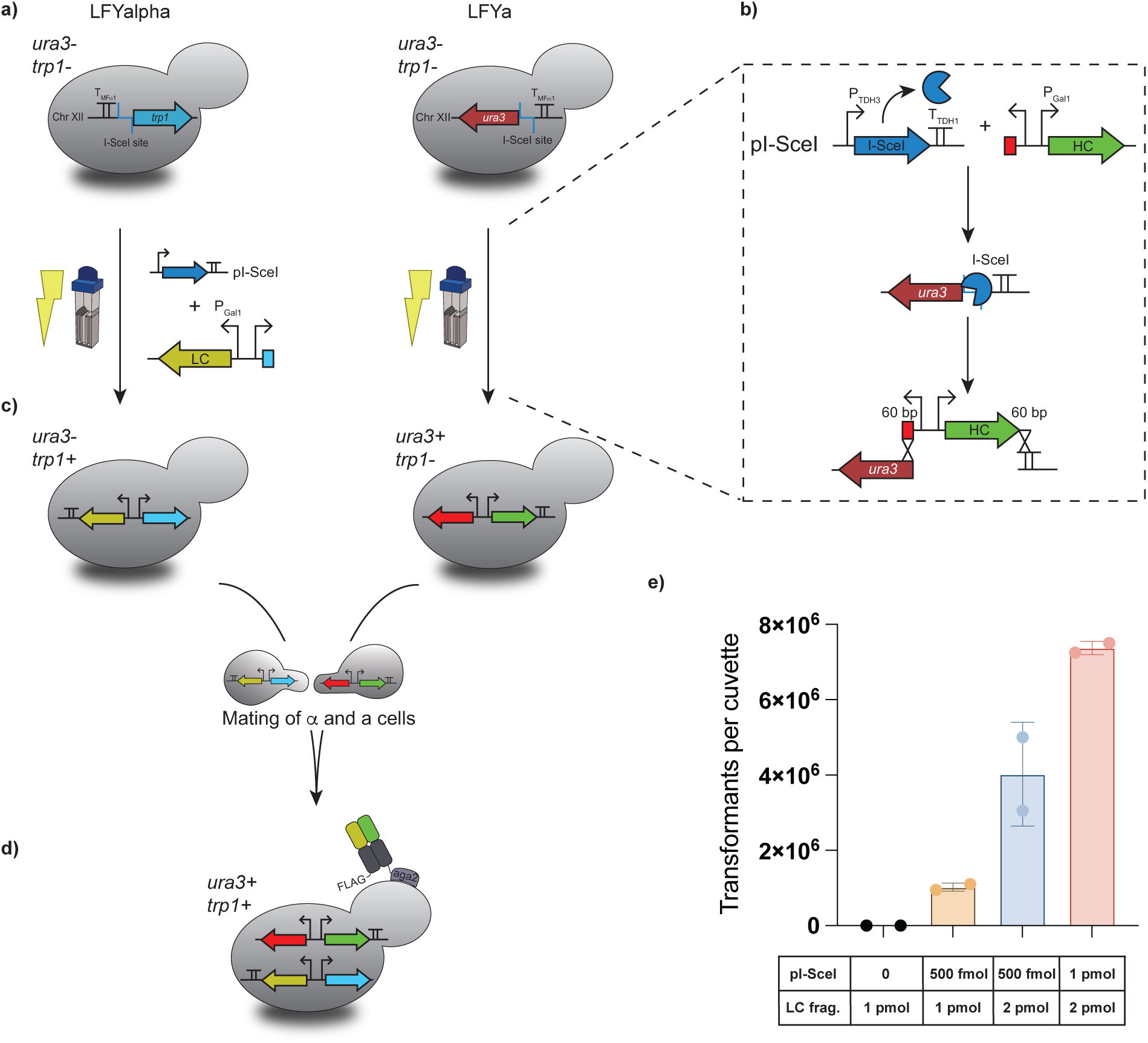
Yeast mating strain design and genomic integration. a) Custom made yeast strains LFYa and LFYalpha carry genomically integrated landing pads on Chr XII. b) Co-transformation of linearized pI-SceI and HC or LC fragments leads to a dsDNA break in the landing pad and thereby facilitates subsequent genomic integration of HC or LC fragments. c) Upon genomic integration, functionality of *trp1* and *ura3* is restored. These auxotrophy markers are used to enrich cells which successfully integrated the HC or LC fragments into the landing pad. d) Mating of MATa and MATα cells leads to MATa/α diploid cells which contain genomically integrated HC and LC and are double positive for *trp1* and *ura3*. e) Different conditions were tested for genomic integration. Varying ratios of pI-SceI and LC fragment (frag.) were experimentally assessed for genomic integration efficiency.

### Experimental screening of yeast mating conditions

For the construction of large combinatorial antibody HC/LC libraries, efficient yeast mating conditions are essential. Previous approaches using yeast mating for the creation of Fab libraries reported mating efficiencies of up to 58% (8). To further optimize mating efficiency, four parameters were systematically tested: (i) carbon sources, (ii) ratio of MATα and MATa cells, (iii) cell density and (iv) mating volume.

To assess mating efficiency of MATα and MATa cells, two spectrally distinct fluorophores, ymNeonGreen and mScarlet-I were genomically integrated into LFYa and LFYalpha, respectively. After mating, the fraction of double positive cells was measured using flow cytometry (**Fig. 2a**). To assess the sensitivity of this method and to verify that double positive events were because of mating, the essential surface protein Sag1 was knocked out in LFYalpha (13). This reduced the mating efficiency from over 80% for functional *sag1* to 0.05%, strongly indicating that the observed double positive events were because of the mating of MATα and MATa cells (**Supplementary Fig. 2a, b**).

**Fig. 2.**
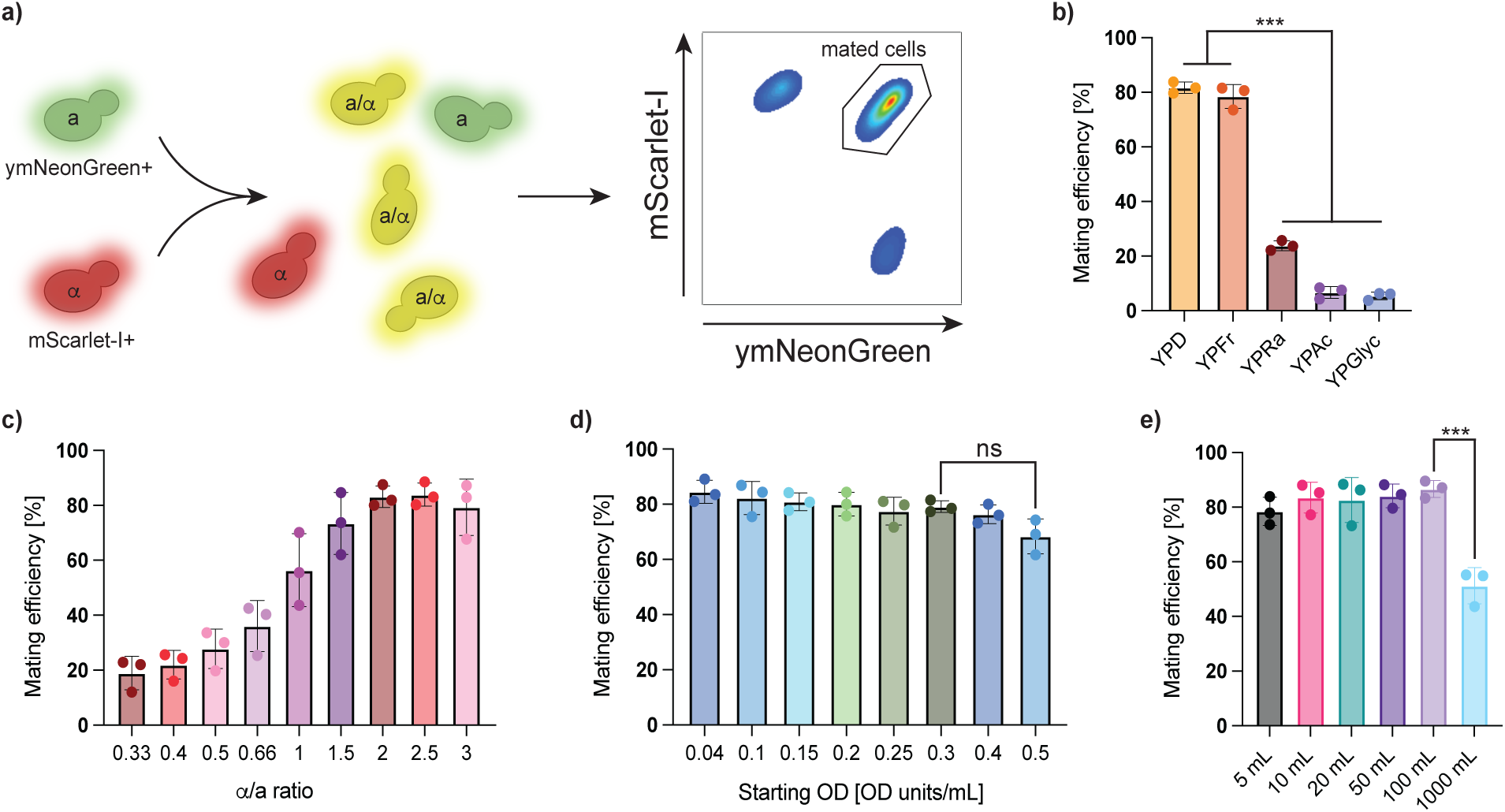
Establishing efficient yeast mating conditions. a) To assess yeast mating efficiency, ymNeonGreen and mScarlet-I were genomically integrated into LFYa and LFYalpha respectively. Mating efficiency is ascertained by quantifying the double positive population using flow cytometry. Mating efficiencies were experimentally tested using b) different carbon sources, c) different ratios of MATα to MATa cells, d) different starting ODs and, e) different culture volumes. (ns; not statistically significant, ***; p-value < 0.01, Methods)

Next, we evaluated different carbon sources and their effect on mating efficiency. Fermentable carbon sources such as glucose (D) and fructose (Fr) led to high mating efficiencies of approximately 80%, a significant increase over non-fermentable carbon sources such as acetate (Ac) or glycerol (Glyc) which yielded low efficiencies of roughly 5%, and raffinose (Ra), a poorly fermentable carbon source, which led to an intermediate efficiency of 20% (**Fig. 2b, Supplementary Fig. 3a**).

Mating efficiency was strongly dependent on the initial ratio of MATα and MATa cells. A 1:1 ratio resulted in an intermediate efficiency of approximately 60%, whereas ratios with fewer MATα to MATa cells produced efficiencies below 20%. Conversely a MATα to MATa ratio of 2.5:1 led to a mating efficiency of approximately 80% (**Fig. 2c, Supplementary Fig. 3b**).

Finally, we determined the impact of initial cell density and culture volume. Mating efficiency remained high at ∼80% for starting optical densities (OD) up to 0.4 units/mL, but dropped below 70%, when the initial OD was increased to 0.5 units/mL. This decrease in mating efficiency was not statistically significant (**Fig. 2d, Supplementary Fig. 3c**). Similarly, scaling the culture volume affected the outcome; a 100 mL volume yielded a high efficiency of 87.5%, whereas increasing the volume to 1, 000 mL significantly reduced the efficiency to ∼55%. (**Fig. 2e, Supplementary Fig. 3d**).

### Testing of recombinases

Previous approaches using yeast mating for shuffled Fab libraries relied on encoding the HC and LC libraries on separate CEN/ARS plasmids (7, 8). Such strategies did not have a mechanism to link HC and LC information together in a single amplicon, and were therefore incompatible with deep sequencing pipelines. Here, we utilize a recombinase-based method to combine the HC/LC information onto one chromosome, enabling efficient deep sequencing. Serine recombinases, BxB1 and ΦBT1 were chosen due to their reported high activity in yeast (19), and were integrated into LFYalpha under the control of the WTC (14). Attachment sites (att) for the recombinases, attP and attB, were placed between HC/LC ORF and T_MFα1_ (**Fig. 3a**) (20). Upon induction with aTc (Methods), the BxB1 recombinase is expressed, recombines the attP and attB sites, thereby creating attL and attR sites and combining HC and LC on the same Chr XII (**Fig. 3a**). The att sites were oriented in such a way that upon recombination, two functioning chromosomes are created. Subsequently, the recombined product can be PCR amplified and deep sequenced. BxB1 and ΦBT1 were tested either with or without C-terminal SV40 nuclear localization signal (NLS). Upon induction, recombination was observed for all designs (**Fig. 3b**). Despite reports of BxB1 being toxic for yeast, it is strictly unidirectional while ΦBT1 has been reported to have bidirectional activity (21, 22). Ultimately, BxB1 without NLS was chosen for subsequent experiments.

**Fig. 3.**
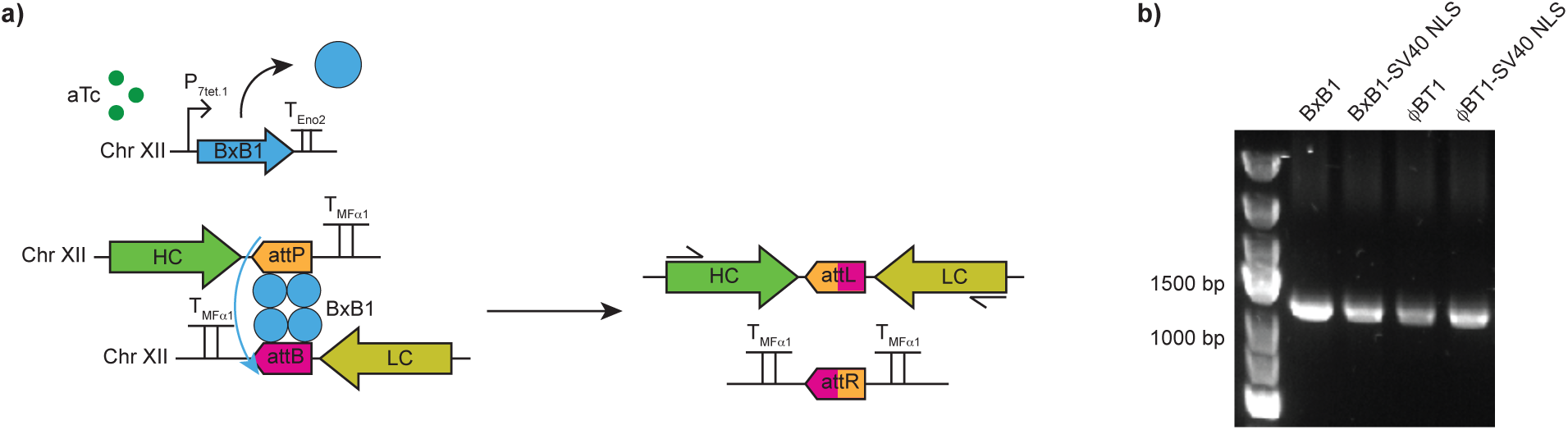
Testing of inducible recombinase. a) Upon addition of aTc, expression of BxB1 recombinase is induced, which subsequently catalyzes the recombination of the two Chr XII, thereby combining HC and LC information onto one chromosome. The combined HC and LC can subsequently be PCR amplified (primers indicated with black arrows) and deep sequenced. b) Testing of BxB1 and ΦBT1 with or without NLS shows that recombination was observed for all conditions.

### Screening of display conditions, library design and construction

To assess and maximize Fab yeast surface display for genomic integration, several parameters were tested. The native aga2 signal sequence and app8, an engineered signal sequence with improved secretion capacity were used (23). Furthermore, we assessed the overexpression of protein disulfide isomerase (PDI), which is an oxidoreductase that is essential for the formation of disulfide bonds in secretory proteins (24). Three different conditions, where PDI was genomically integrated in LFYa and expressed using P_RNR2_, P_HHF1_ and P_TDH3_ (weak, medium and strong constitutive promoter) were tested (18). When using the app8 signal sequence for the HC, a significant increase in surface display was observed when compared to the aga2 signal sequence (**Fig. 4a**, Methods). Notably, this increase in surface display level was independent of the signal sequence used for the LC. When testing different levels of PDI overexpression, no further gain in surface display level was observed (**Fig. 4a**). Based on this observation, the app8 signal sequence was chosen for the HC library, while the aga2 signal sequence was utilized for the LC library.

**Fig. 4.**
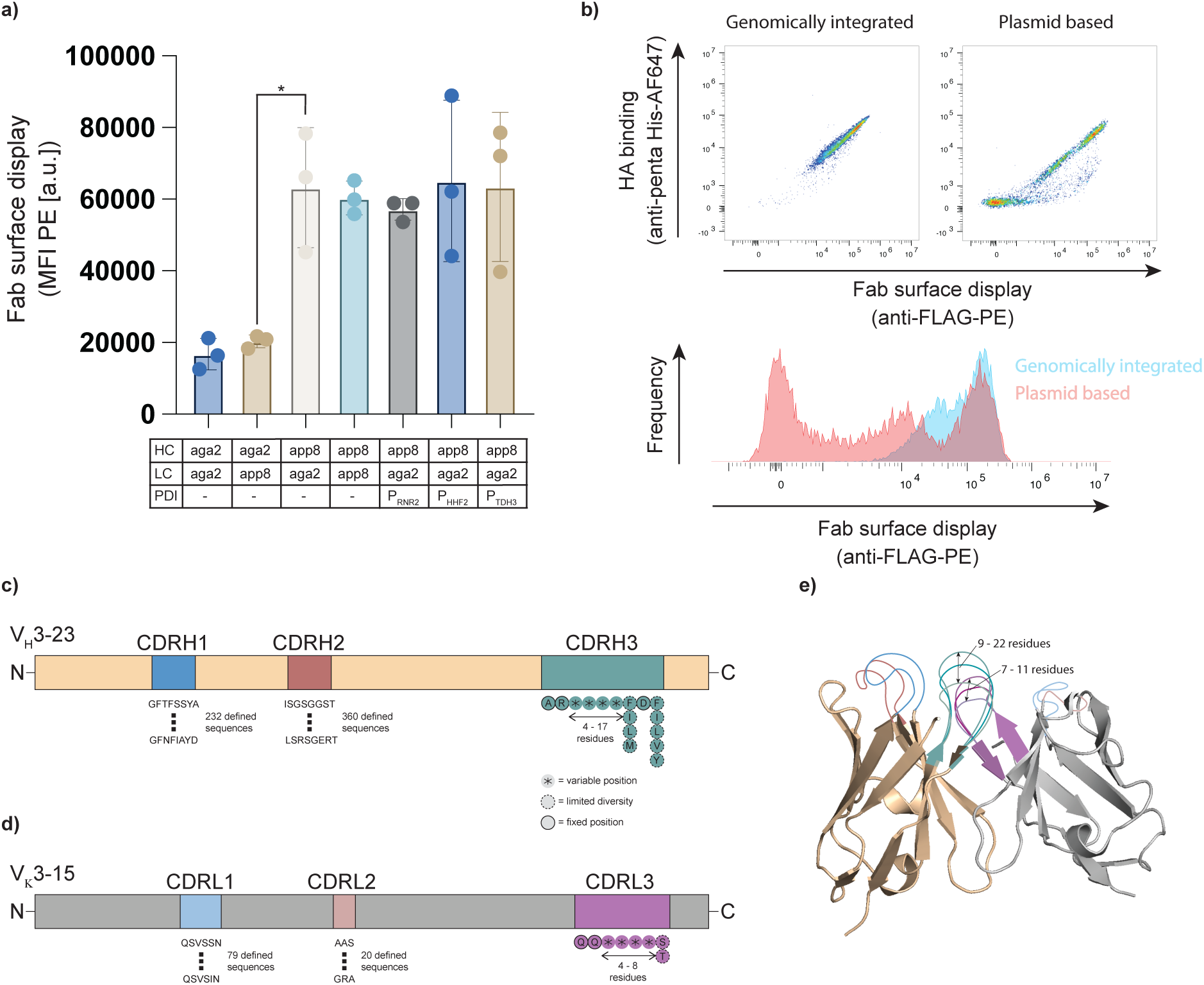
Testing of surface display conditions and synthetic antibody library design. a) Different combinations of signal sequences and PDI expression levels were tested to maximize Fab surface display. MFI; median fluorescence intensity, (*; p-value < 0.05, Methods). b) Dot plots for genomically integrated (top-left) versus plasmid based (top-right) Fab surface display. Bottom: Overlayed histograms for Fab surface display for genomically integrated (blue) and plasmid based (red) expression. c) Schematic of synthetic HC library. The framework regions (FR) in beige were kept constant, while the CDR loops were varied. d) Schematic of synthetic LC library. The FRs in gray were kept constant, while the CDR loops were varied. e) Structure of the combined HC and LC libraries.

To understand the impact of genomic integration on display, plasmid-encoded versus genomically integrated HC/LC was experimentally investigated. For this purpose, we used a model antibody (clone H2214 with specificity to Influenza HA antigen) with the germline background of V_H_3-23 for HC and V_Κ_3-15 for LC (25). Expression with two separate plasmids resulted in more than a third of all cells not expressing measurable levels of Fab surface display. Conversely, nearly all cells that had HC and LC genomically integrated show Fab surface display. Furthermore, plasmid-encoded HC/LC information led to large display heterogeneity with display levels spanning five orders of magnitude, whereas genomic integration demonstrated more homogeneous display levels (**Fig. 4b**).

Furthermore, for the display of monomeric proteins, we engineered the strain LFY100 based on LFYa through integration of the functional *trp1* gene (**Supplementary table 1**). Following library integration, this allows the cells to grow in minimal medium without the need for mating. When comparing the surface display of a genomically integrated affibody Z_HER2:342_ in LFY100 and a plasmid encoded control, a more homogenous display was observed for the genomically integrated variant. Nearly all cells with genomically integrated affibody exhibiting measurable levels of surface display, while for the plasmid encoded condition, more than a third of all cells showed no display (**Supplementary Fig. 4**).

Next, we designed a synthetic antibody library based on V_H_3-23*01 and V_K_3-15*01 human germlines for HC and LC, respectively. This pairing has previously been described as having advantageous biophysical properties (26). For the HC, J_H_4 and for the LC, J_Κ_2 was chosen due to their frequent use in nature (27, 28). Next, we analyzed approximately 36, 000 HC and 11, 000 LC sequences from IMGT. In order to maximize functionality, the diversity for CDRH1, CDRL1, CDRH2, and CDRL2 was limited to observed sequences. 232 sequences for CDRH1, 360 sequences for CDRH2, 79 sequences for CDRL1 and 20 sequences for CDRL2 were utilized for the library construction. For CDRH3 and CDRL3, recombined sequences were analyzed, and degenerate codons were designed to mimic natural amino acid (aa) frequencies (29). Certain positions show low natural diversity, for example positions 1 and 2 of CDRH3 strongly favor Ala and Arg respectively, thus these were fixed (**Fig. 4c, d**). The natural length diversity was approximated by covering CDRH3 lengths of 9 - 22 aa and CDRL3 lengths of 7 - 11 aa (**Fig. 4c-e**).

Libraries for CDRH1, CDRL1, CDRH2 and CDRL2 were ordered as oligonucleotide pools, while for the CDRH3 and CDRL3 libraries, degenerate primers were ordered (**Supplementary Table 2**) for each length and subsequently pooled in ratios to mimic their natural length distribution. For both HC and LC, the covered lengths comprise more than 90% of the natural length diversity. The libraries were then assembled using overlap extension PCR (OE-PCR) and genomically integrated, yielding library sizes of 4.1×10^7^ for the HC and 1.7×10^7^ for LC (**Supplementary Fig. 5, Supplementary Table 3**, Methods). The individual HC and LC libraries were mated and based on experimentally determined mating efficiencies and culture volume, the library diversity was estimated to be ∼1.2×10^11^ (Methods). The libraries were deep sequenced pre- and post-mating. The length distribution of HC and LC library both pre-and post-mating was in close agreement with the designed length distribution (**Supplementary Fig. 6a, b**). Furthermore, both libraries pre- and post-mating showed aa diversities that closely matched the designed diversities (**Supplementary Fig. 6c, d**).

### Antibody discovery from yeast mating libraries

Post-mating, expression staining showed that approximately 94% of cells had measurable Fab surface display (**Supplementary Fig. 7**), which far exceeds previously reported rates of 20% (8). Next, we screened our antibody library for binding against several protein antigen targets: human tumor-necrosis factor-alpha (TNFα), human epidermal growth factor receptor 2 (HER2) and Influenza hemagglutinin (HA) (Darwin variant). In an initial step, the library was selected using one round of magnetic-associated cell sorting (MACS) followed by five rounds of FACS with decreasing antigen concentrations (**Supplementary Fig. 8**, Methods). After the final round of selection using 500 pM antigen, BxB1 expression was induced, and the sorted populations were prepared for deep sequencing using the PacBio platform (Methods). For TNFα, one primary clone (CDRH3-CDRL3) pairing dominated after the selection, making up more than 90% of the population (**Fig. 5a, Supplementary table 4**). Screening against HER2 resulted in several clones showing strong enrichment, with HC sequences paired in combination with a range of different LC sequences (**Fig. 5a, Supplementary table 5**). For HA, one highly enriched HC variant was identified in combination with a variety of LC sequences (**Fig. 5a, Supplementary table 6**).

**Fig. 5.**
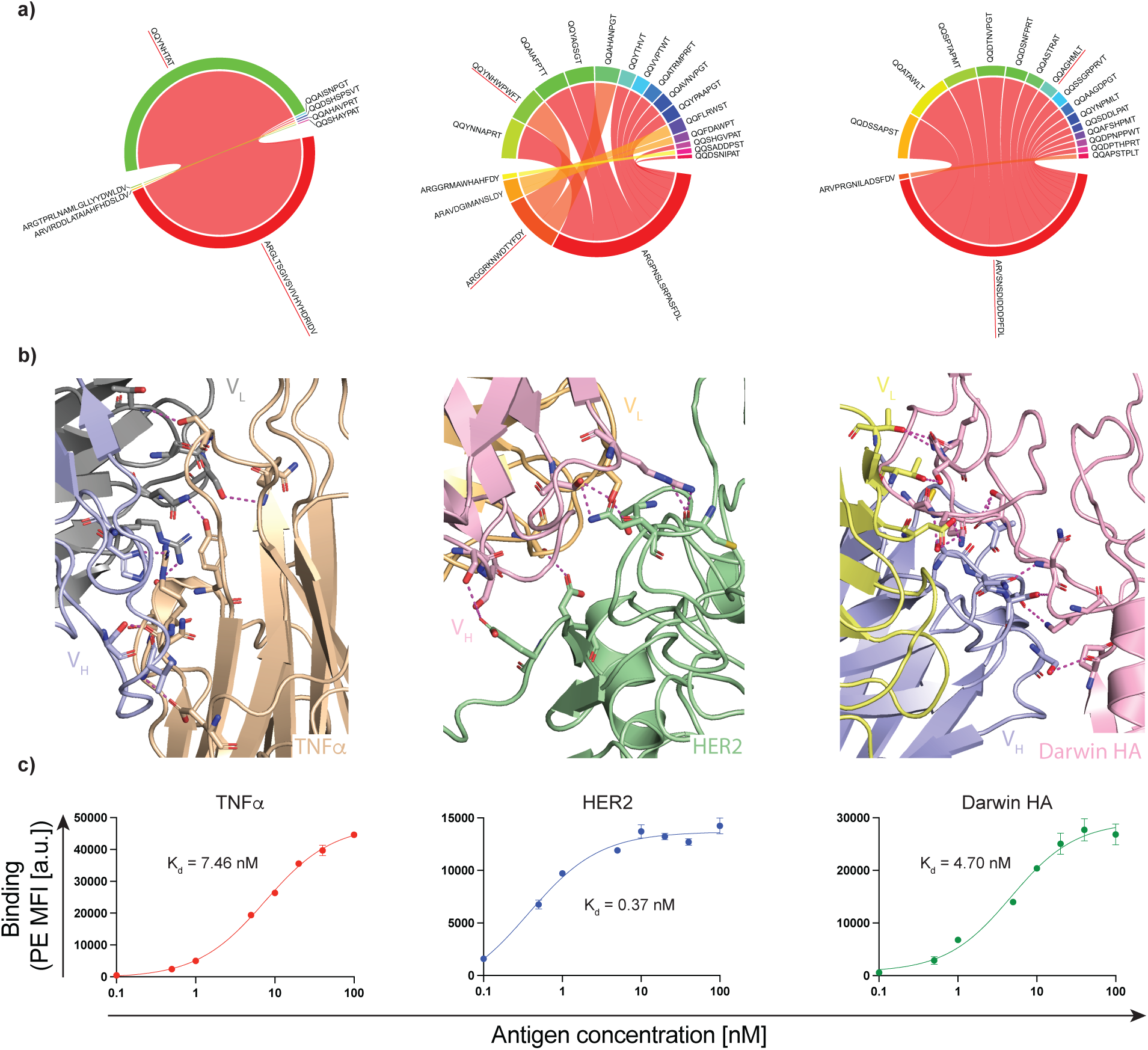
Characterization of antibody binders to target antigens. a) Circos plots depicting the association of CDRH3 (bottom half) and CDRL3 (top half) sequences for TNFα (left), HER2 (middle) and HA (right). The top 5 most frequent variants for TNFα and top 15 most frequent variants for HER2 and HA are shown. Variants underlined with red were further characterized. b) AlphaFold3 structural predictions for binding interactions of selected antibodies to cognate antigen. c) Affinity predictions measured by yeast surface display of Fab, antigen titration and flow cytometry.

The most abundant antibody targeting TNFα was predicted by AlphaFold3 to bind near the core β-sandwich structure (30) with a K_d_ of 7.46 nM. Similarly, one of the top variants for HER2 and HA were predicted to bind domains III and IV (31) and the stem region (32), exhibiting high affinities with K_d_ values of 0.37 nM and 4.70 nM, respectively (**Fig. 5b, c, Supplementary Fig. 9a-c, Supplementary Fig. 10a-c, Supplementary Table 7**).

## Discussion

While YSD has become a cornerstone of antibody engineering, its utility has been constrained by two persistent challenges: limited library diversity compared to phage display, and heterogeneous surface expression caused by plasmid-based systems. Our work addresses these limitations through an integrated platform combining genomic integration, optimized yeast mating, and recombinase-mediated linkage of HC and LC information. The resulting system achieves phage-scale library diversity (>10^11^ variants) while maintaining the eukaryotic advantages of yeast display.

A key innovation of this study is the development of LFYa and LFYalpha strains featuring genomic landing pads for high-efficiency, stable library integration. Unlike plasmid-based systems where more than a third of cells lose Fab expression (**Fig. 4b**), genomically integrated libraries show uniform surface display without decrease over time. Nearly 95% of the mated library showed measurable surface display, compared to 20% in previous studies using yeast mating for Fab engineering (8). This homogeneity is critical for quantitative screening, as expression variability can obscure affinity during FACS. This consistent surface expression level may enable more accurate high-throughput quantitative measurements of apparent affinity using titration-based screening and deep sequencing (33).

The optimized mating protocol developed here represents another substantial advance. By identifying glucose as the optimal carbon source and establishing a 2.5:1 MATα:MATa ratio, we achieved 87.5% mating efficiency, a 30% improvement over previous reports (8). This high efficiency ensures that library diversity is preserved during the mating process, a crucial factor when working with rare clones in large libraries. We hypothesize that the observed differences in mating efficiency with regards to MATα:MATa ratio are because of the influence of the mating pheromones. a-and α-factors arrest the cell cycle. If a-factor inhibits the growth of MATα cells more strongly than vice versa, it would explain why more MATα cells per MATa cell are required to achieve efficient mating.

The incorporation of BxB1 recombinase addresses a critical bottleneck in yeast mating approaches; the inability to link HC/LC sequences for deep sequencing. By combining both chains onto a single chromosome (**Fig. 3a**), our platform enables direct sequencing without requiring cumbersome single-cell PCR or subcloning. This enabled us to isolate antibodies with therapeutically relevant K_d_ values in the single-digit nanomolar to picomolar affinity range for a variety of antigens.

Looking forward, this system opens several research avenues. The strains could be used for the engineering of dimeric therapeutic proteins beyond Fab, such as T-cell receptors. We also envision LFY100 as a valuable platform for the engineering of monomeric scaffolds, like nanobodies, DARPins, and affibodies (34, 35), providing an alternative to EBY100 for more homogeneous surface display.

In summary, we have developed a yeast display platform that bridges the gap between prokaryotic-scale diversity and eukaryotic protein processing. By solving long-standing issues of library size, stability, and compatibility with deep sequencing, this work expands the toolbox for antibody discovery and protein engineering.

## Methods

### Genomic modifications

BJ5464 and BJ5465 were engineered using the MyLO protocol (12). Briefly, yeast cells were grown overnight in YPD (20 g/L vegetable peptone (Millipore, 18332), 10 g/L yeast extract (Millipore, 70161), 20 g/L glucose (Sigma-Aldrich, G8270)). On the day of the genomic modification, the cells were diluted to an OD_600_ of 0.3 in 50 mL YPD and incubated at 30 °C and 250 RPM until an OD_600_ of 1.6 was reached. The cells were conditioned by directly adding 0.5 mL 1M Tris/1M DTT (Roche, 10708984001) and 2.5 mL 2M LiAc·2H_2_O/TE (Sigma-Aldrich, L6883). The cells were incubated for 15 min at 30 °C. The cells were washed three times using Electroporation Buffer (1 M Sorbitol (Sigma-Aldrich, S6021), 10 mM Trizma base (Sigma-Aldrich, T1503) and 1 mM calcium chloride (Merck, 1.02382.0500), pH 7.5). The electrocompetent cells were mixed with 100 fmol linearized plasmid carrying a guide RNA (gRNA), constitutively expressed SpCas9 and an antibiotic resistance marker and 200 fmol HDR template. gRNA sequences are reported in **Supplementary Table 8**. The HDR templates were flanked by 70 - 200 bp homology to the targeted sequence in the genome. The cells were electroporated using 2 mm gap cuvettes (Sigma-Aldrich, Z706086) and 2.49 kV using a Bio-Rad MicroPulser. The cells were recovered for 2 h in YPD at 30 °C before plating them on YPD agar plates with antibiotics (G418 (Invivogen, ant-gn-5) or Hygromycin (Invivogen, ant-hg-5), 200 μg/mL final concentration) and incubated at 30 °C for 3 days. To verify the genomic modifications, single colonies were picked, lysed for 1 h at 37 °C using Zymolyase (Zymo, E1004) and subsequently used as template for yeast colony PCR (cPCR). For yeast cPCR, PhirePlant polymerase was used (Thermo Fisher, F-160). For primer sequences for the individual loci, see **Supplementary table 9**.

### Screening of mating conditions

Fluorescent LFYa and LFYalpha were grown separately overnight in sterile filtered YPD at 30 °C and 250 RPM to stationary phase. For different carbon sources, the following recipes were used: YPD (10 g/L yeast extract, 20 g/L peptone, 20 g/L glucose), YPFr (10 g/L yeast extract, 20 g/L peptone, 20 g/L fructose (Sigma-Aldrich, F0127)), YPRa (10 g/L yeast extract, 20 g/L peptone, 20 g/L raffinose (Roth, 5241.5)), YPAc (20 g/L yeast extract, 20 g/L peptone, 10 g/L KOAc (Sigma-Aldrich, 60035)) and YPGlyc (10 g/L yeast extract, 20 g/L peptone, 30 mL/L glycerol (Roth, 7530.1)). All media was sterile filtered using a 0.22 μm filter. For mating volumes less than 20 mL, round bottom bacterial cultivation tubes were used. For volumes of 20 mL or larger, non-baffled Erlenmeyer flasks were used. The flasks were filled 20% of the maximum volume. The cells were mated for 24 h at 30 °C in a shaking incubator set to 250 RPM. 10 μL of cells were washed using 200 μL autoMACS running buffer (Miltenyi, 130-091-221), resuspended using 200 μL autoMACS running buffer and run on a Cytoflex. 10000 events were recorded for each condition.

### Library construction using OE-PCR

For the construction of HC and LC libraries, three different plasmids were prepared. The first construct contained the partial auxotrophy marker, P_ura3_ for HC or P_trp1_ for LC, P_Gal1_, and FR1. The second construct contained only FR3 and the third construct contained FR4, C_H_1 or C_L_, aga2 for HC and FLAG tag for LC and the partial T_MFα1_ (**Supplementary Fig. 5a**). The rationale for putting the individual parts onto different plasmids is to prevent template amplification during the OE-PCR. The three plasmids were used for the construction of the separate CDR1, 2 and 3 libraries using PCR (Q5 HotStart, NEB, M0494) (**Supplementary Fig. 5b**). Afterwards, 5 μL ExoI (NEB, M0293) was directly added to the PCR and incubated for 1 h at 37 °C to remove any remaining primers. The PCR products were column purified (Zymo, C1003). In a subsequent step, using primer pairs T_MFα1_ fwd + ura3 rev for HC and T_MFα1_ fwd + trp1 rev for LC (**Supplementary table 9**), the individual fragments were assembled into a single piece using OE-PCR (**Supplementary Fig. 5c**). For the OE-PCR, 50 fmol of each fragment was used per PCR. For the annealing temperature, 68 - 69 °C was used. The assembled libraries were column purified (Zymo, C1019) and directly used for transformation in yeast.

### Library transformation

LFYa and LFYalpha cells were grown overnight in YPD. On the day of library transformation, the cells were diluted to an OD_600_ of 0.3 in a total volume of 300 mL YPD + 10x AA (1.688 g/L Yeast Synthetic Drop-out Medium Supplements (Sigma-Aldrich, Y2001), 0.022 g/L uracil (Sigma-Aldrich, U0750), 0.209 g/L L-histidine monohydrochloride monohydrate (Sigma-Aldrich, H8125) and 0.082 g/L L-tryptophan (Sigma-Aldrich, T8941)). The cells were incubated at 30 °C until an OD_600_ of 1.6 was reached. The cells were conditioned by directly adding 3 mL 1M Tris/1M DTT and 15 mL 2M LiAc·2H_2_O/TE. The cells were incubated for 15 min at 30 °C. The cells were washed three times using Electroporation Buffer. 95 μL competent cells were mixed with 1 pmol linearized pI-SceI and 2 pmol insert (DNA was concentrated to 5 μL). pI-SceI can be linearized with either a restriction digest using BamHI or PCR amplified using the primer pair pI-SceI fwd and pI-SceI rev (**Supplementary table 9**). In this work, pI-SceI was linearized using PCR. The cells were filled into a 2 mm gap cuvette and electroporated with 1.5 kV using a Bio-Rad MicroPulser. Afterwards, the cells were recovered in 200 mL YPD + 10x AA for 1h at 30 °C. The cells were pelleted and transferred to 500 mL selective SD-CAA medium (20 g/L glucose, 0.856 g/L NaH_2_PO_4_·H_2_O (Roth, K300.1), 0.677 g/L Na_2_HPO_4_·2H_2_O (Sigma-Aldrich, 1.06580), 6.7 g/L yeast nitrogen base without amino acids (Sigma-Aldrich, Y0626) and 5 g/L casamino acids (Gibco, 223120) in deionized water). For LFYa, SD-CAA has to be further supplemented with 2 mM L-tryptophan (Sigma-Aldrich, T0254) and for LFYalpha with 1.33 mM uracil. The cells were incubated at 30 °C for 24 h. 500 OD_600_ units were transferred into 500 mL fresh selective SD-CAA medium and incubated for another 24 h at 30 °C. Transformation efficiency was determined using dilution plating.

### Mating of libraries

To mate the separate libraries, they were first grown separately overnight to stationary phase. Subsequently, the cells were pooled in 100 mL sterile filtered YPD for a starting cell density of 0.3 OD_600_ units/mL and a MATα:MATa ratio of 2.5:1. The cells were incubated in a 500 mL non-baffled Erlenmeyer flask for 24 h at 30 °C. After mating the cells, 1000 OD_600_ units were pelleted and transferred into 1000 mL SD-CAA and incubated for 24 h at 30 °C. The cells were passaged a second time by pelleting 1000 OD_600_ units and transferring the cells into 1000 mL SD-CAA. The cells were incubated for another 24 h at 30 °C.

### Screening of Fab library

Libraries grown in SD-CAA were diluted into 1 L SG-CAA (20 g/L galactose (Sigma-Aldrich, G0625), 0.856 g/L NaH_2_PO_4_·H_2_O, 0.677 g/L Na_2_HPO_4_·2H_2_O, 6.7 g/L yeast nitrogen base without amino acids and 5 g/L casamino acids in deionized water) for a starting cell density of 1 OD_600_ unit/mL. The cells were incubated at 23 °C and 250 RPM for 48 h. In an initial selection step, the cells were selected using MACS. 8×10^9^ cells were pelleted and washed using 40 mL cold autoMACS running buffer. Subsequently, the cells were stained using 100 nM antigen in 40 mL for 1 h at 4 °C. The cells were pelleted and washed using 40 mL cold autoMACS running buffer. The cells were resuspended using 38 mL cold autoMACS running buffer and 2 mL Streptavidin MicroBeads (Miltenyi, 130-048-101). The cells were incubated for 30 min at 4 °C. The cells were pelleted and resuspended using 12 mL cold autoMACS running buffer. 4 LS columns (Miltenyi, 130-042-401) were equilibrated using 3 mL cold autoMACS rinsing solution (Miltenyi, 130-091-222). 3 mL cell suspension was loaded per column and run through a 30 μm strainer (Miltenyi, 130-041-407). The column was washed three times with 5 mL cold autoMACS running buffer. The cells were eluted from the column using 5 mL cold autoMACS running buffer. The eluted cells were pelleted and resuspended using 10 mL SD-CAA. The cells were incubated for 2 days at 30 °C.

Subsequently, the cells were screened using FACS. Streptavidin binders were depleted before every round of sorting. 5×10^7^ cells were washed using 1 mL cold autoMACS running buffer and resuspended using 900 μL cold autoMACS running buffer and 100 μL Streptavidin MicroBeads. The suspension was incubated at 4 °C for 30 min on a rotor. The cells were pelleted and resuspended using 1 mL cold autoMACS running buffer. LD columns (Miltenyi, 130-042-901) were equilibrated using 3 mL cold autoMACS running buffer. The cells were loaded on the LD column and the column was washed twice using 1 mL cold autoMACS running buffer. The flow-through was collected and pelleted. The collected cells were subsequently stained with antigen for 1 h at 4 °C and 700 RPM. The staining volume was adjusted accordingly to maintain a 10 fold excess of antigen. The cells were subsequently washed using 1 mL cold autoMACS running buffer and stained with 1 mL Streptavidin-PE (BioLegend, 405204) and anti-FLAG-APC (BioLegend, 637307) staining solution. Streptavidin-PE was diluted 1:100 and anti-FLAG-APC 1:200 using cold autoMACS running buffer. The cells were incubated for 30 min at 4 °C and 700 RPM. The cells were washed using 1 mL cold autoMACS running buffer. The cells were resuspended using 2 mL cold autoMACS running buffer and sorted on an AriaFusion.

### Induction of Bxb1 expression, genomic DNA extraction and deep sequencing

Cells were grown overnight at 30 °C and 250 RPM in YPD to stationary phase. The cells were subsequently passaged into 20 mL fresh YPD + 1 μM aTc (Cayman Chemical, Cay10009542) (10 mM aTc stock solution in DMSO (Sigma-Aldrich, D4540)) at a density of 0.25 OD units/mL. The cells were incubated in a 100 mL round-bottom Erlenmeyer flask for 24 h at 30 °C and 250 RPM. Subsequently, genomic DNA was extracted from the cells using the Yeast Genomic DNA kit from Thermo Fisher (Thermo Fisher, 78870). Briefly, 7 mL of culture was pelleted, yielding a cell pellet of roughly 100 mg in weight. The pellet was resuspended using 8 μL/mg pellet Y-PER buffer. The cells were incubated at 65°C for 10 min. The cells were pelleted for 5 min at 13000 g. The supernatant was discarded and the pellet was resuspended using 400 μL of each DNA Releasing Reagent A and B. The cells were incubated at 65°C for 10 min. 200 μL of Protein Removal Reagent was added and the tube was inverted 10 times. The cells were pelleted for 5 min at 13000 g. The supernatant was transferred to a new 2 mL Eppendorf tube and 600 μL isopropanol was added. The tube was inverted 10 times and subsequently centrifuged for 10 min at 13000 g. The supernatant was discarded and 1.5 mL 70% ethanol was added to the tube. The tube was inverted 10 times and centrifuged for 1 min at 13000 g. Residual ethanol was removed by drying the sample in a vacuum centrifuge. The DNA was solubilized using 50 μL nuclease-free water. The libraries were PCR amplified using primers binding the signal sequence (**Supplementary table 9**). The libraries were sequenced using either a MiSeq i100 for separate HC and LC libraries or a PacBio Revio for joined HC/LC libraries.

### Statistical analysis

For all statistical analysis, a two-sided t-test assuming unequal variance was applied.

### Material availability

pI-SceI and integration vectors for LFYa and LFYalpha (pLFYa and pLFYalpha) were deposited on Addgene (Addgene ID 243802 - 243804). LFYa, LFYalpha and LFY100 were deposited on ATCC.

## Acknowledgements

We thank Marco Mandolesi and Joseph Taft for helpful discussions and input. We thank William Kelton and Kai-Lin Hong for valuable input and suggestions on the manuscript. We thank the ETH Zurich Department of Biosystems Science and Engineering Single Cell Unit and the Functional Genomics Center Zurich for their support.

## Contributions

L.F. conceived the study and developed the methodology. L.F. designed and executed all experiments and performed data analysis. L.F. and S.T.R. wrote the manuscript.

## Supplementary tables

**Supplementary Table 1.** Genomic modifications for LFY100.

| Engineered strains | Parent strains | Introduced genomic modifications |
| --- | --- | --- |
| LFY100 | BJ5465 | *ura3-Δ*, *trp1*, *sag1::P<sub>Gal1</sub>* *aga1*, XII-2::landing pad a |

**Supplementary Table 2.**
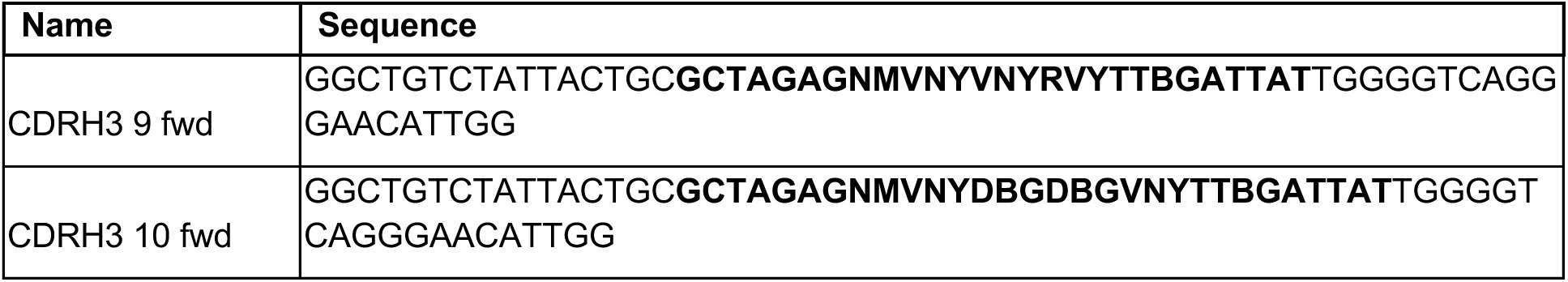
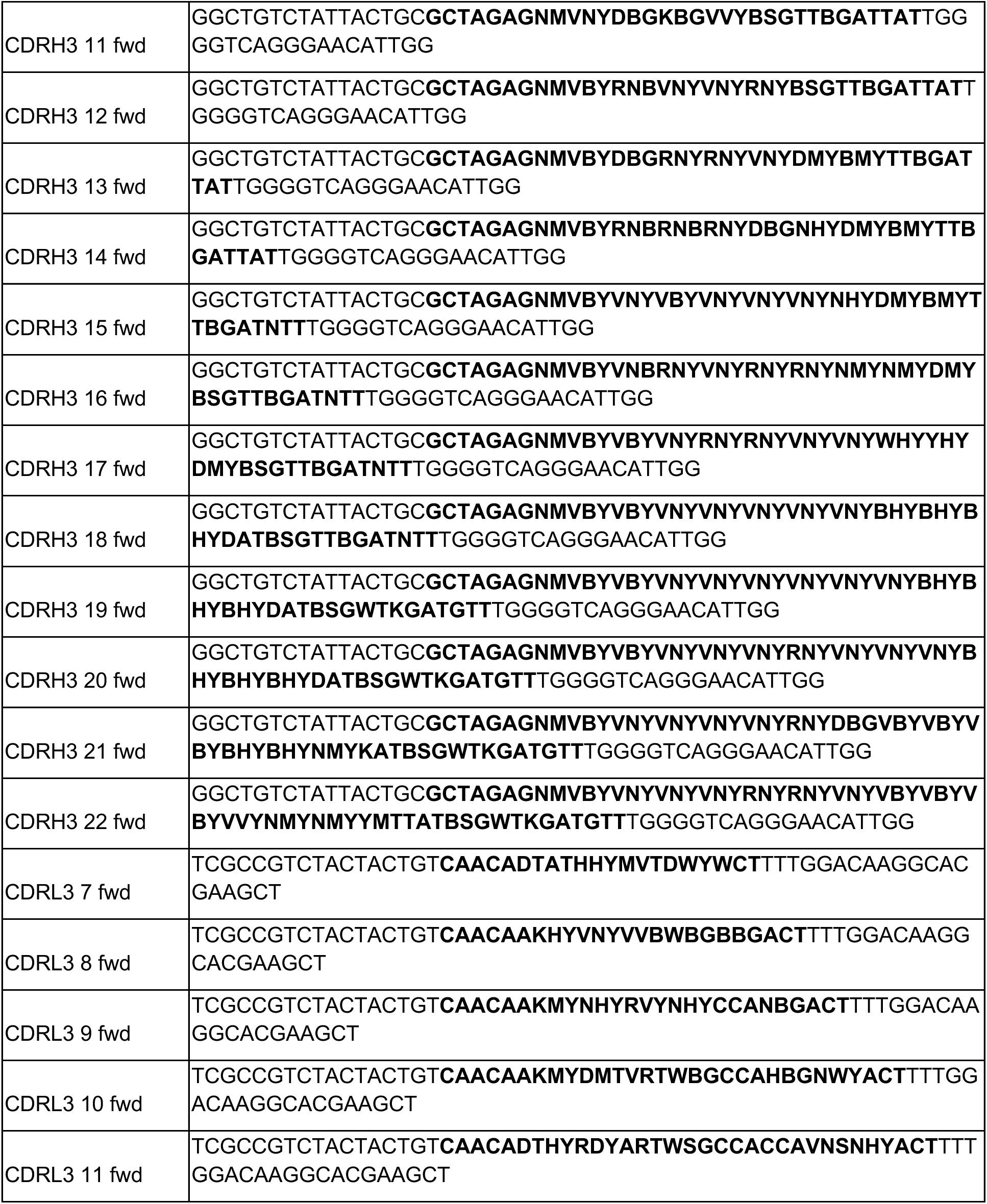
Degenerate codon design for CDRH3 and CDRL3. Sequences highlighted in bold indicate the CDR3 regions.

**Supplementary Table 3.** Library sizes for HC and LC

|  |  |
| --- | --- |
|  | Library size |
| HC library | $4.1 \times 10^7$ |
| LC library | $1.7 \times 10^7$ |

**Supplementary Table 4.**
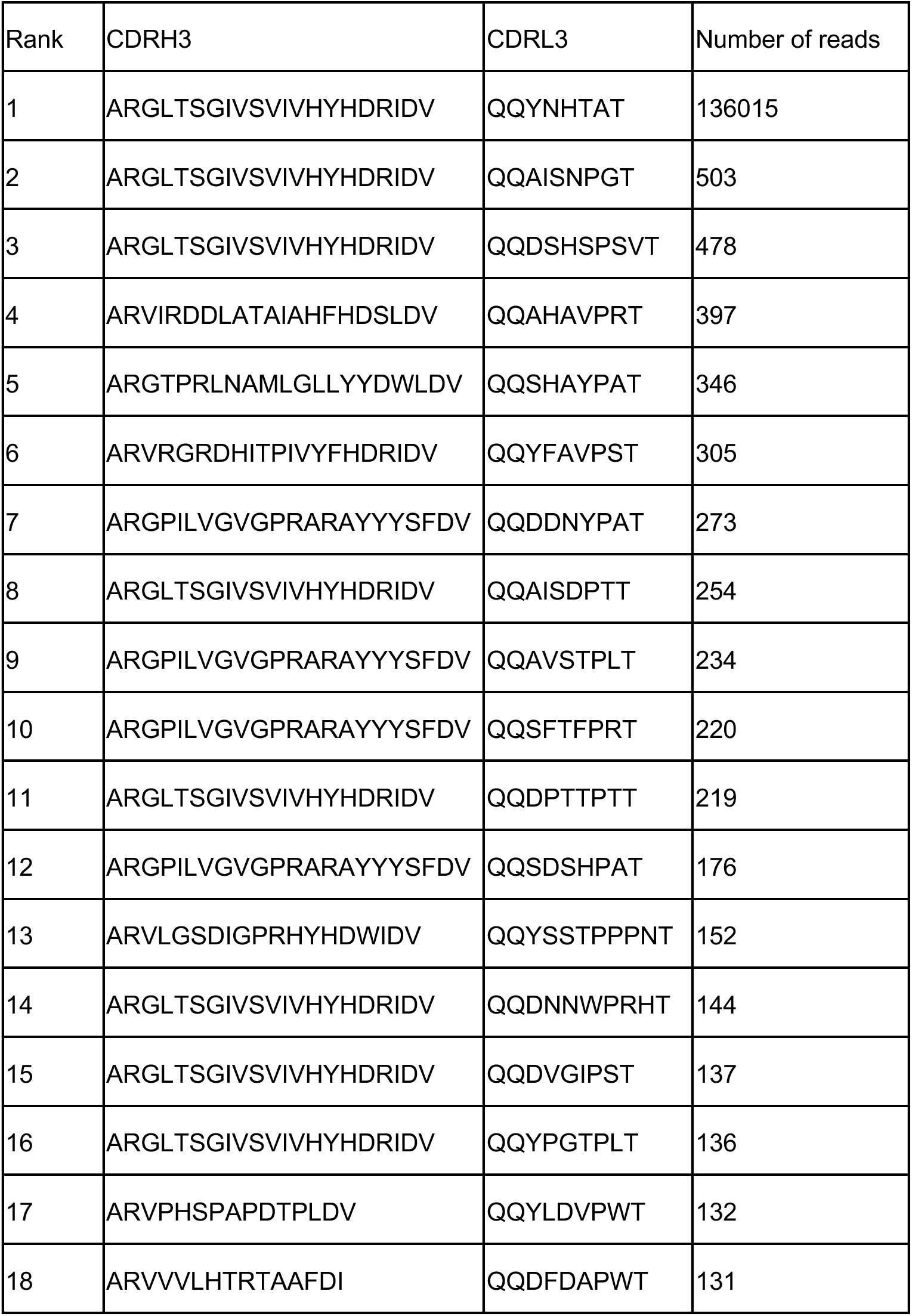
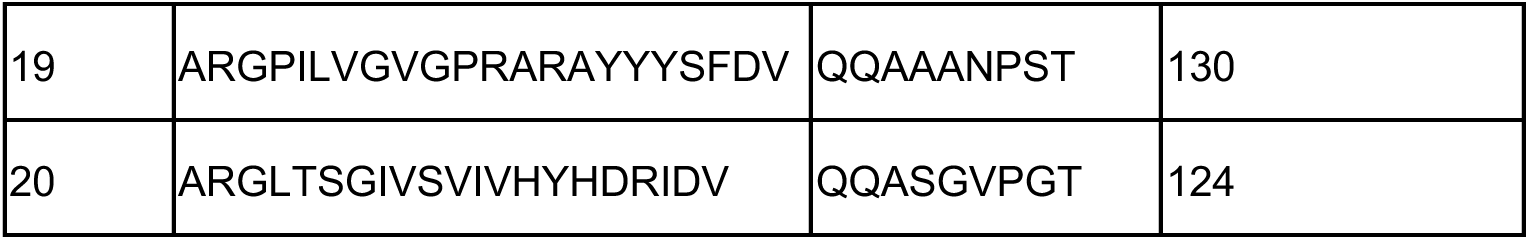
Ranked sequences by number of reads of CDRH3 and CDRL3 for TNFα binders.

**Supplementary Table 5.**
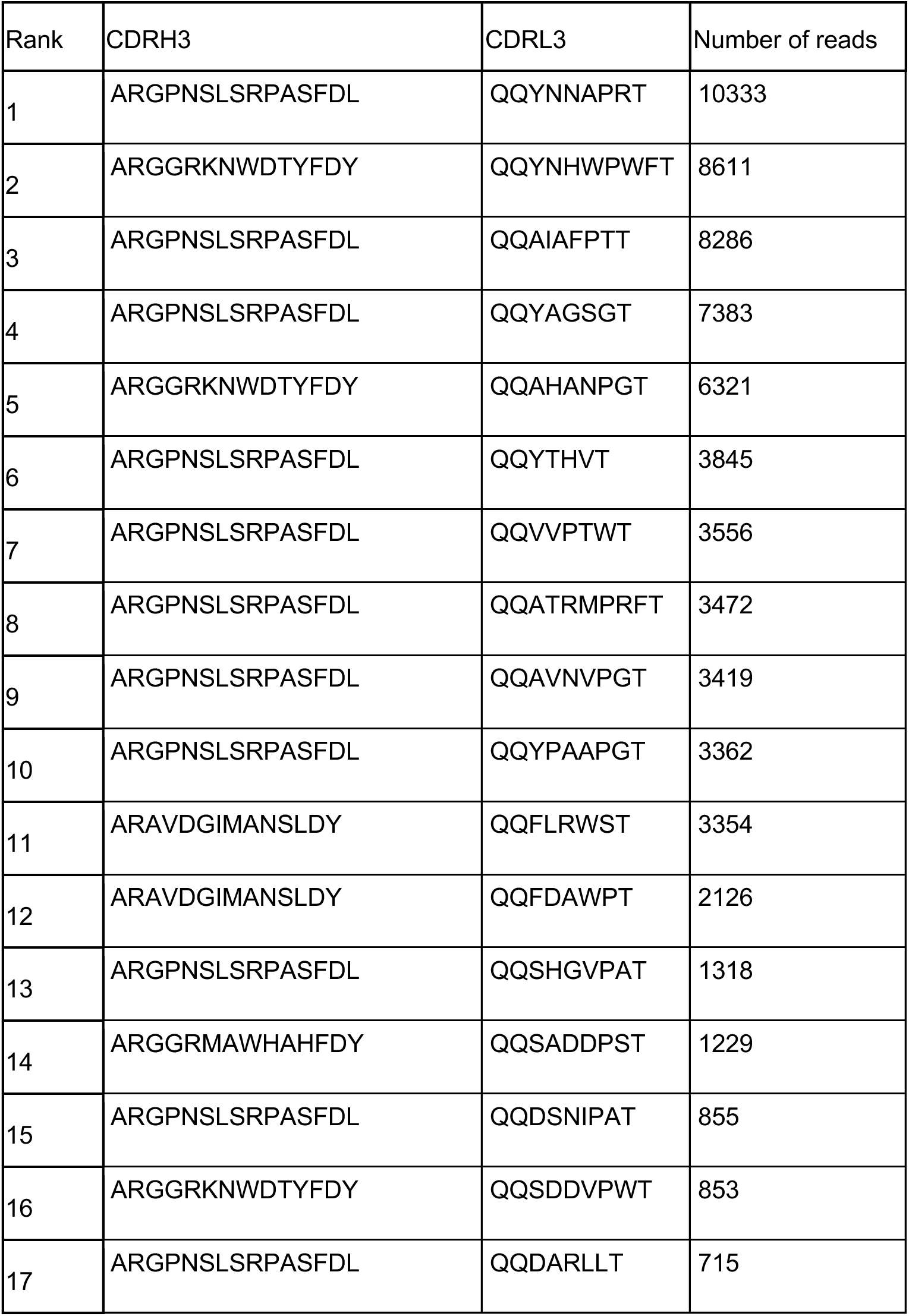
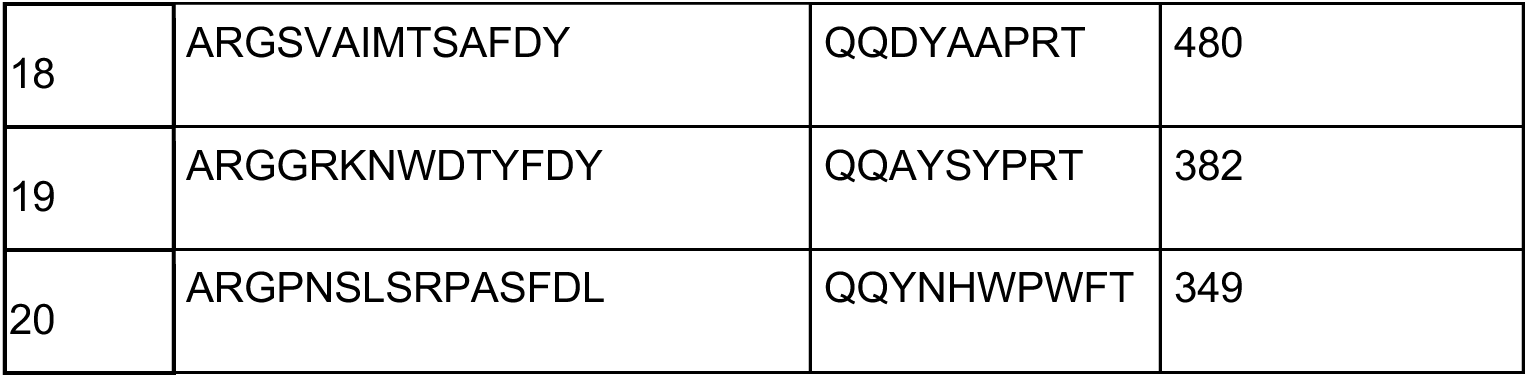
Ranked sequences by number of reads of CDRH3 and CDRL3 for HER2 binders.

**Supplementary Table 6.**
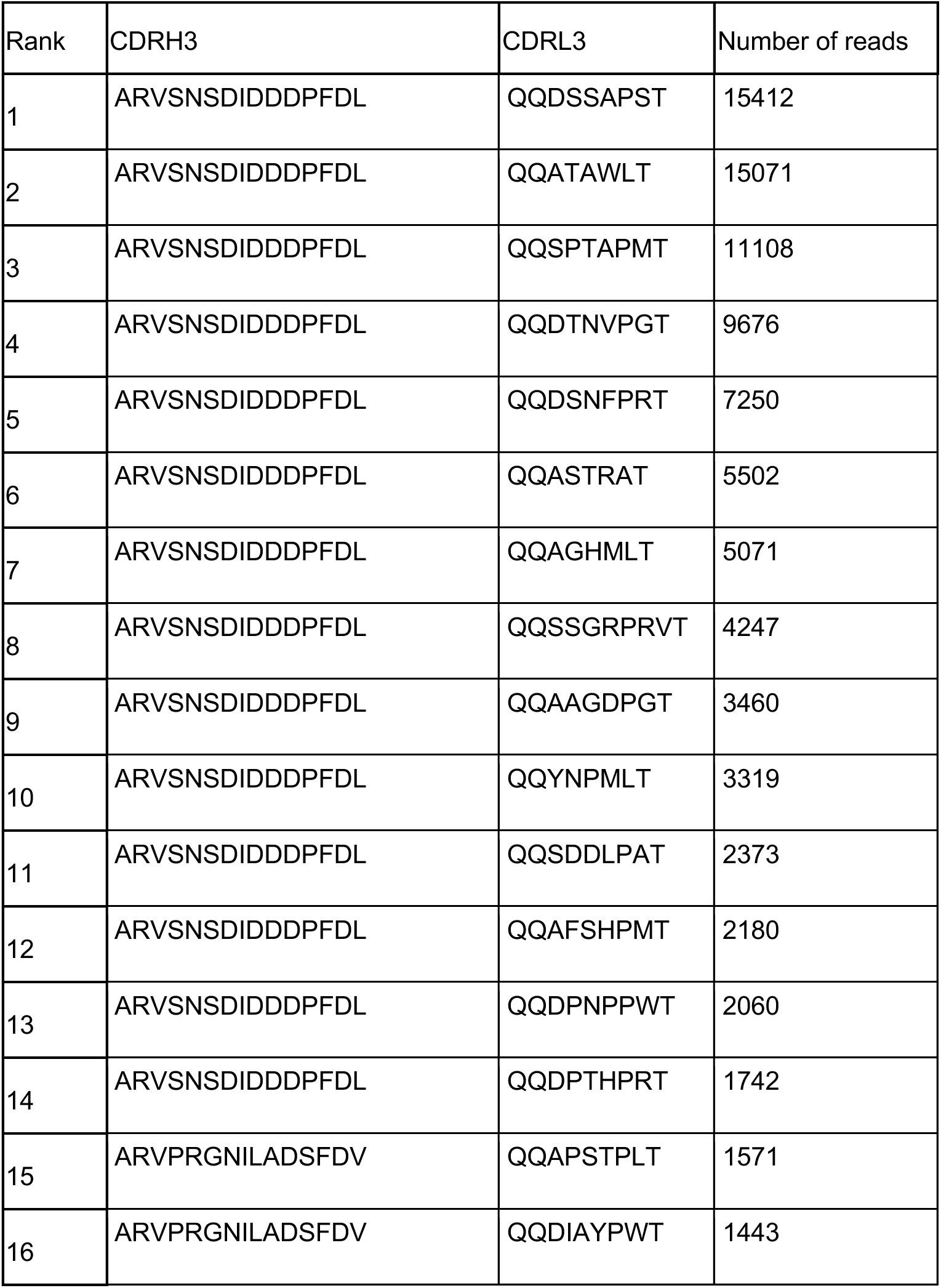
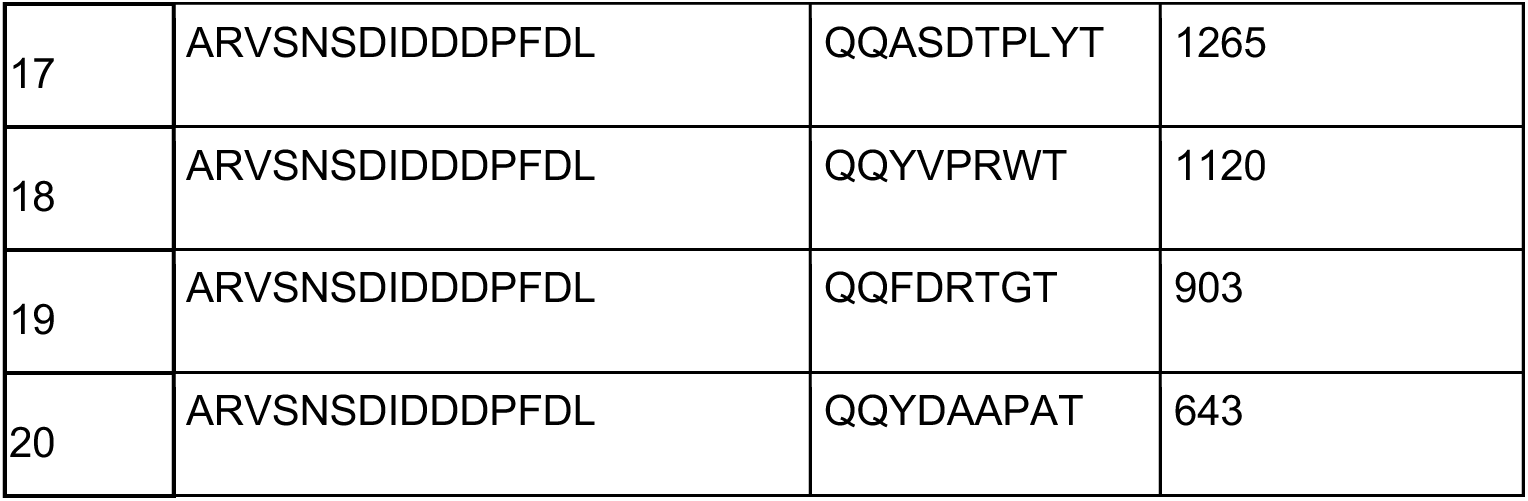
Ranked sequences by number of reads of CDRH3 and CDRL3 for Darwin HA binders.

**Supplementary Table 7.**
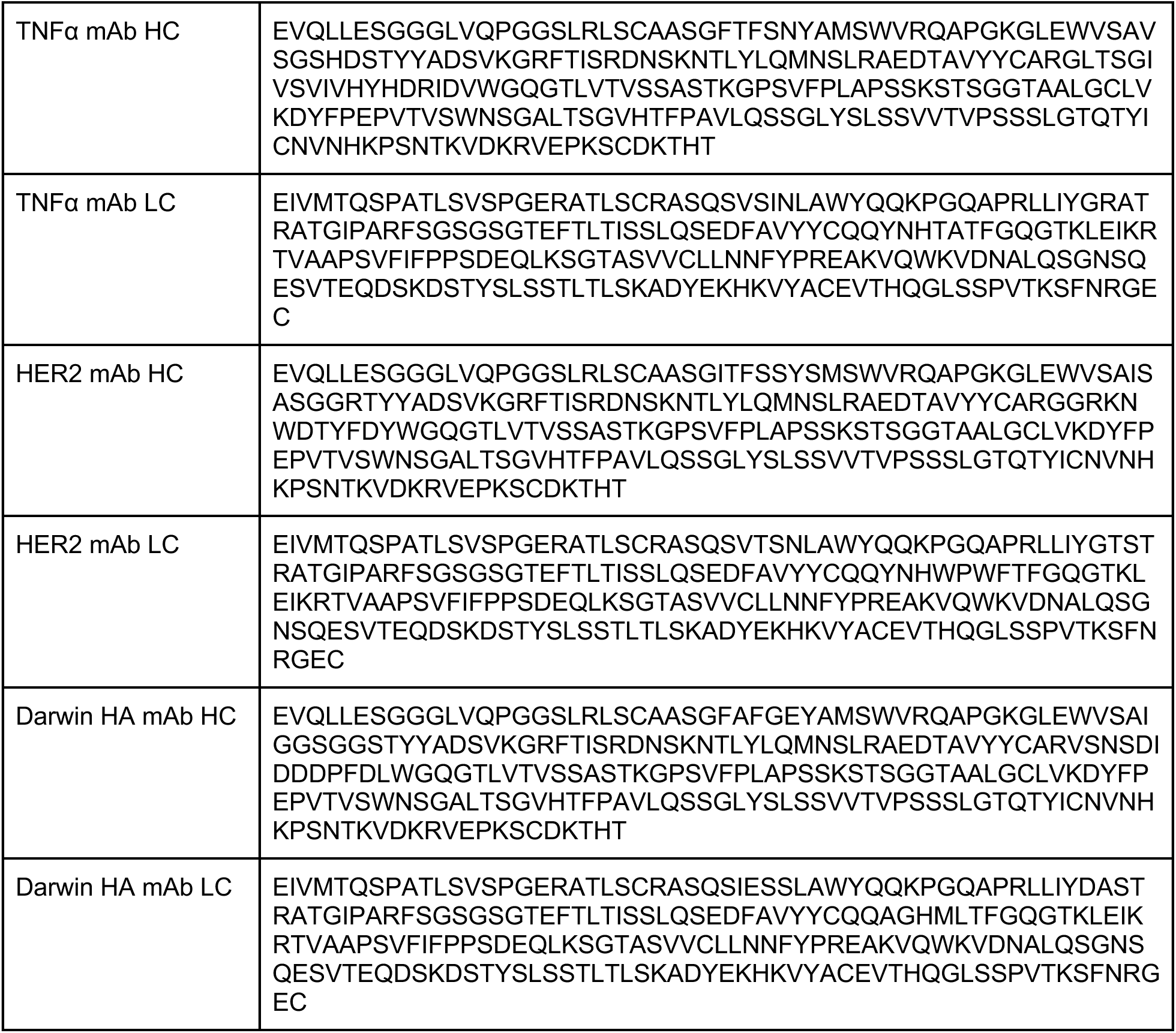
HC and LC sequences for tested antibodies.

**Supplementary Table 8.**
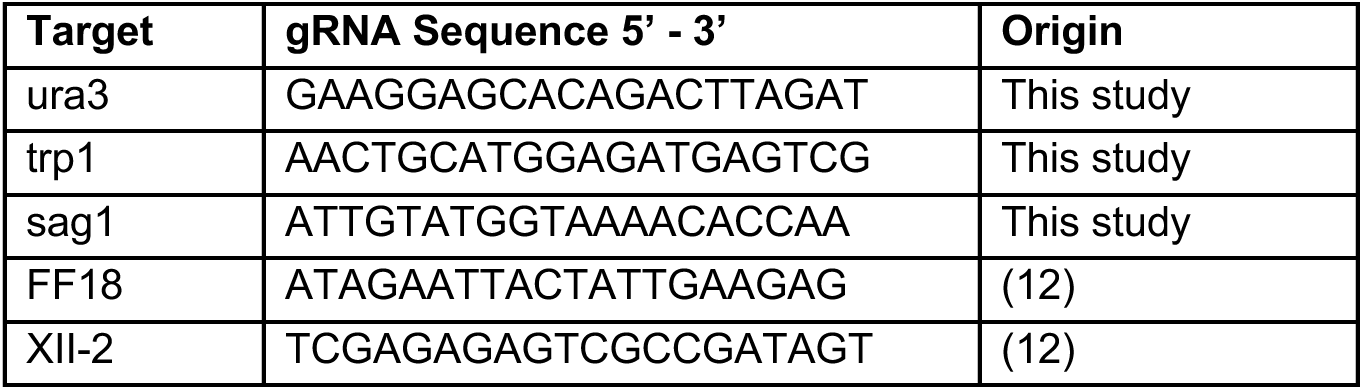
gRNAs used in this study to introduce genomic modifications.

**Supplementary Table 9.**
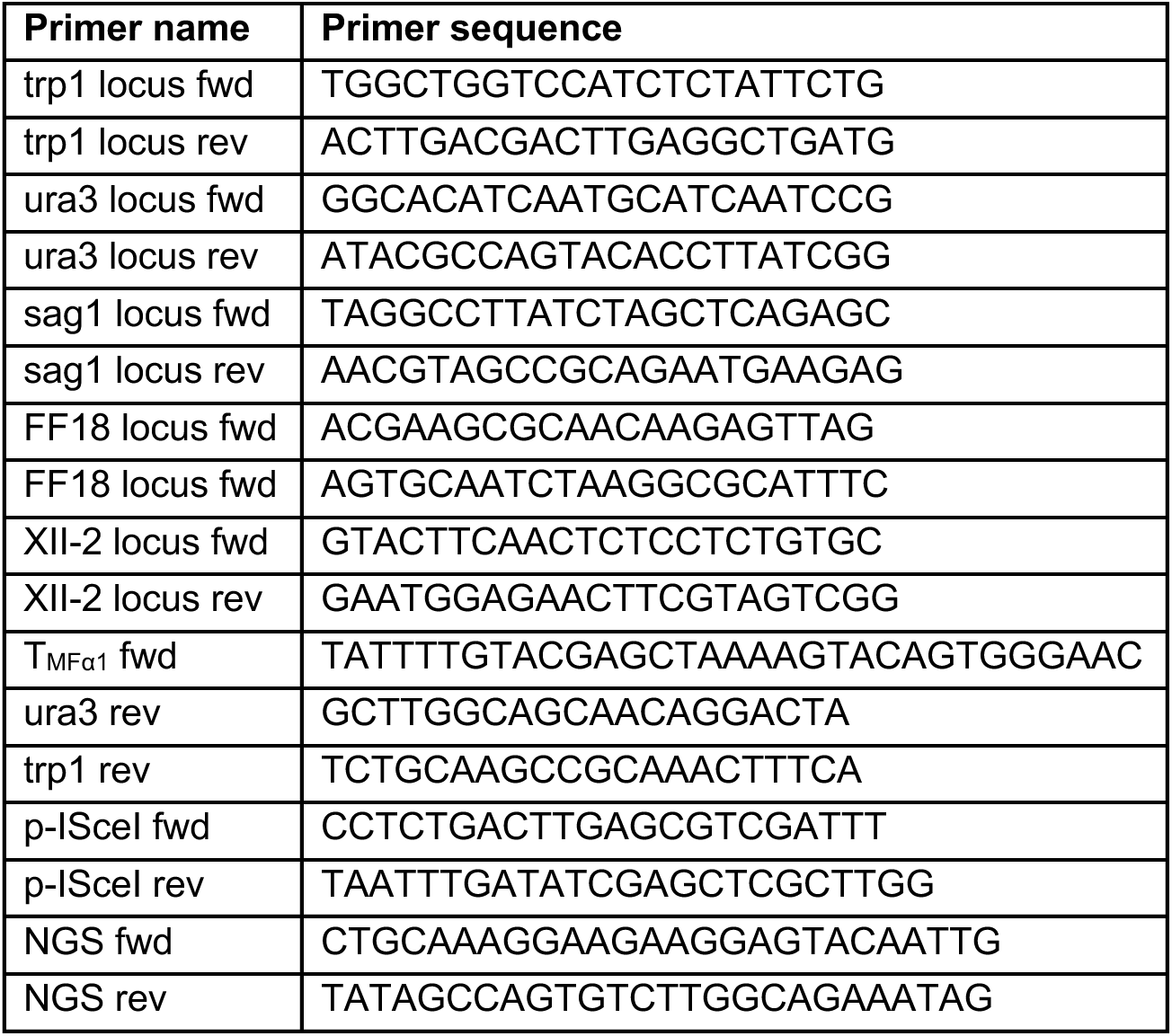
Primers used in this study.

## Supplementary figures

**Supplementary Fig. 1.** Screening of different conditions for genomic integration. Dilution plating for the four tested conditions. For the condition 0 pI-SceI and 1 pmol LC frag, a 1:10^3^ dilution is shown. For the remaining conditions, 1:10^5^ dilutions are shown. A 1:10^5^ dilution means that every observed colony equals 10^5^ transformation events. On every plate, the transformation result from two cuvettes was plated.

**Supplementary Fig. 2.**
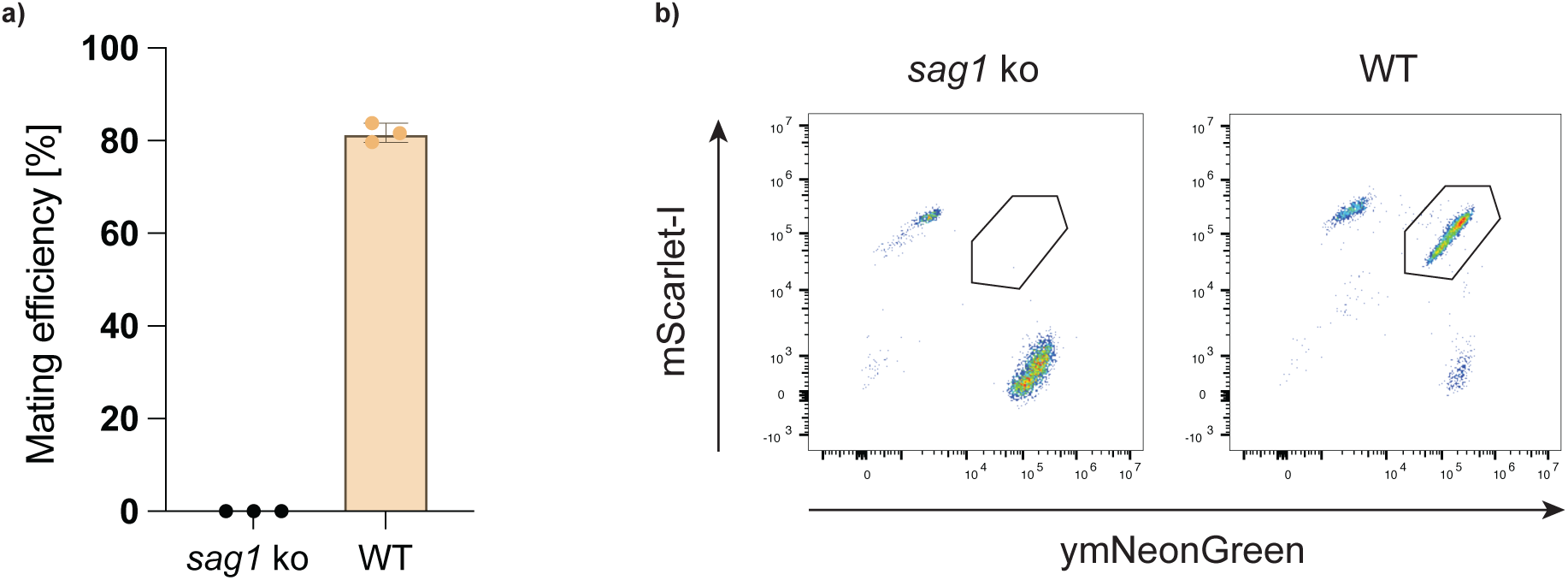
Mating efficiency without and with functional *sag1* in LFYalpha. Mating efficiency between LFYa and LFYalpha with a *sag1* knock out (ko) or functional *sag1*. Mating efficiency is represented as a) bar plot and b) dot plot.

**Supplementary Fig. 3.**
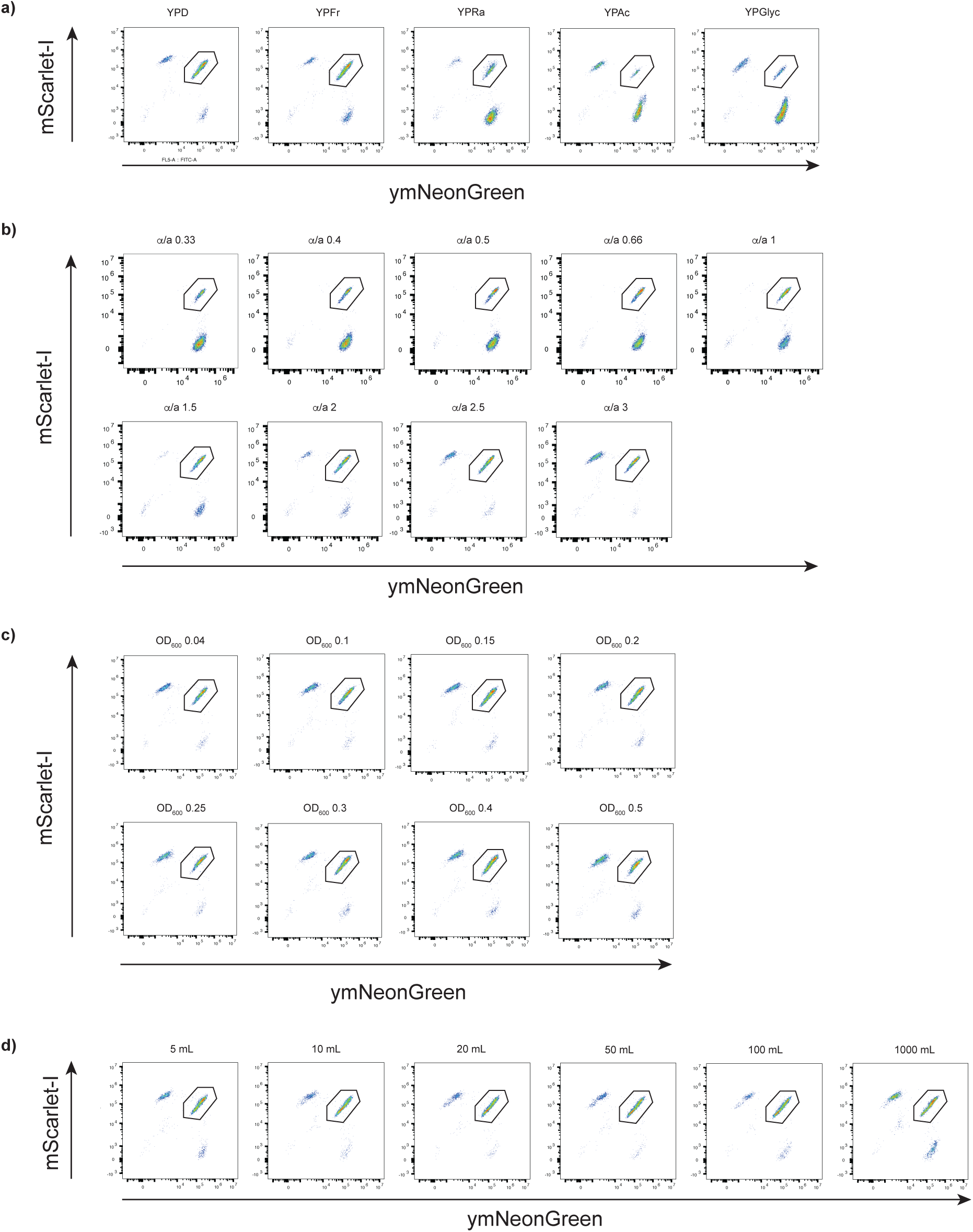
Dot plots from mating experiment. a) Different carbon sources. b) Different ratios of MATα and MATa cells. c) Different starting ODs. d) Different mating volumes.

**Supplementary Fig. 4.**
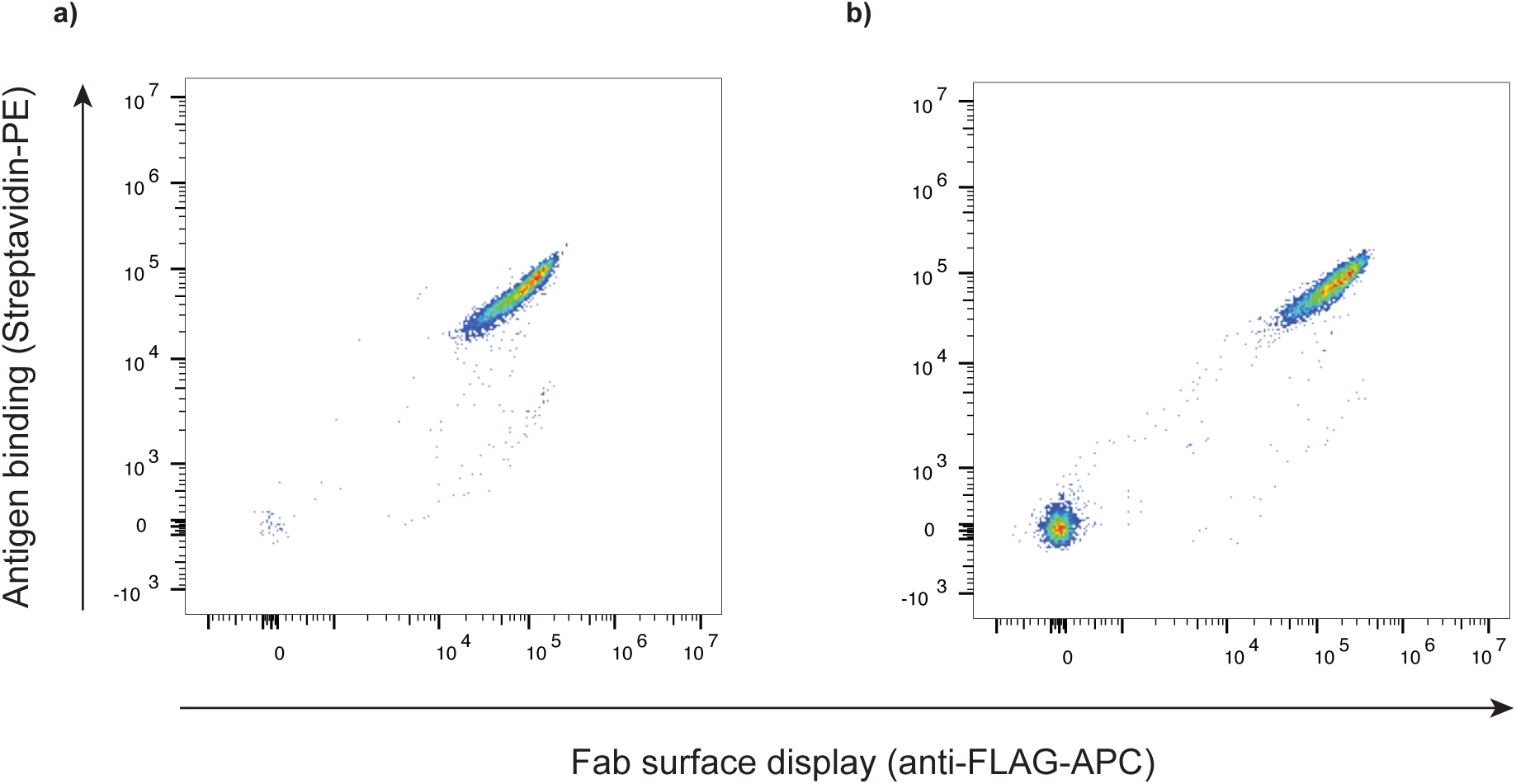
Surface display of monomeric protein in LFY100. Surface display of ZHER2:342, a) genomically integrated into LFY100 and b) encoded on a plasmid.

**Supplementary Fig. 5.**
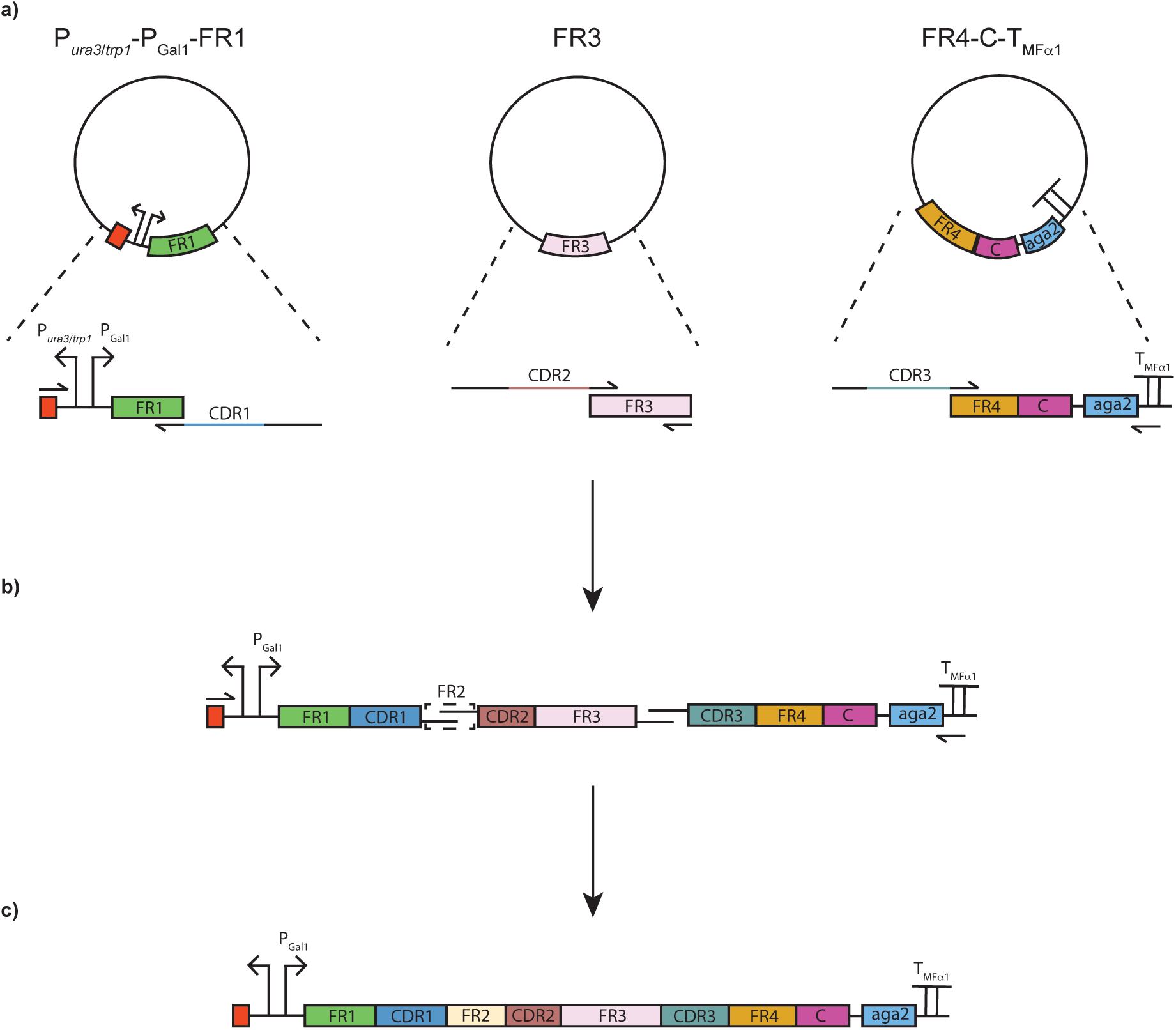
Library construction using OE-PCR. a) For both HC and LC libraries, three plasmids were prepared, carrying P*ura3*/*trp1*-PGal1-FR1, FR3 and FR4-C-TMFα1. For the HC library, the third plasmid contains the sequence for aga2, while for the LC library, this plasmid carries the FLAG tag sequence. The plasmids are used as templates for PCRs during which the CDR1, 2 and 3 diversity is introduced. C; constant region of HC or LC. b) For OE-PCR, the three fragments were pooled in equimolar ratios and primers binding to ura3/trp1 and TMFα1 were utilized to link the fragments together. c) The resulting product from the OE-PCR was directly used for transformation into LFYa and LFYalpha.

**Supplementary Fig. 6.**
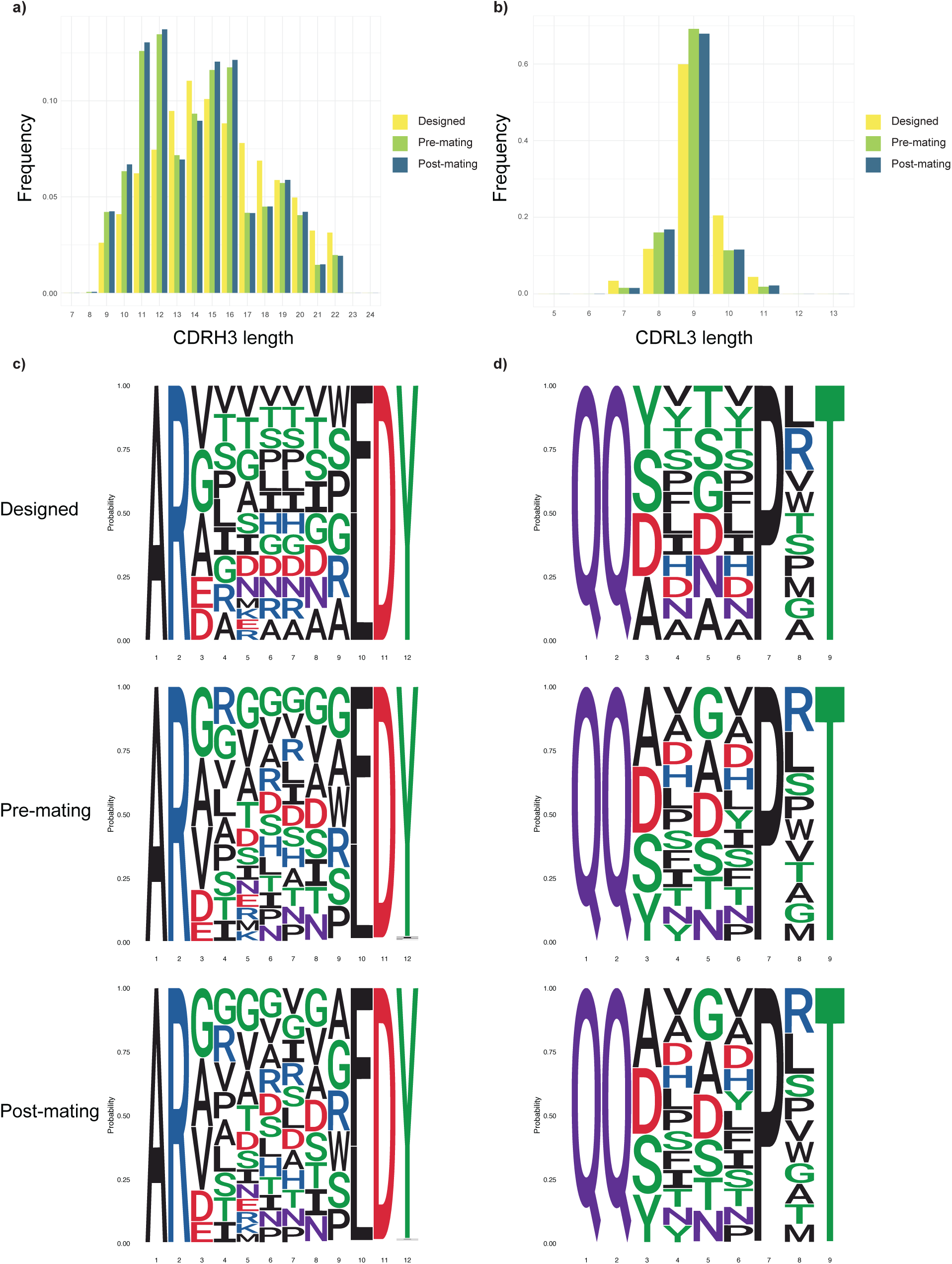
LC and HC library characterization. Length distribution for a) CDRH3 and b) CDRL3. Sequence logo plot for designed, pre- and post-mating libraries for c) CDRH3 and d) CDRL3.

**Supplementary Fig. 7.**
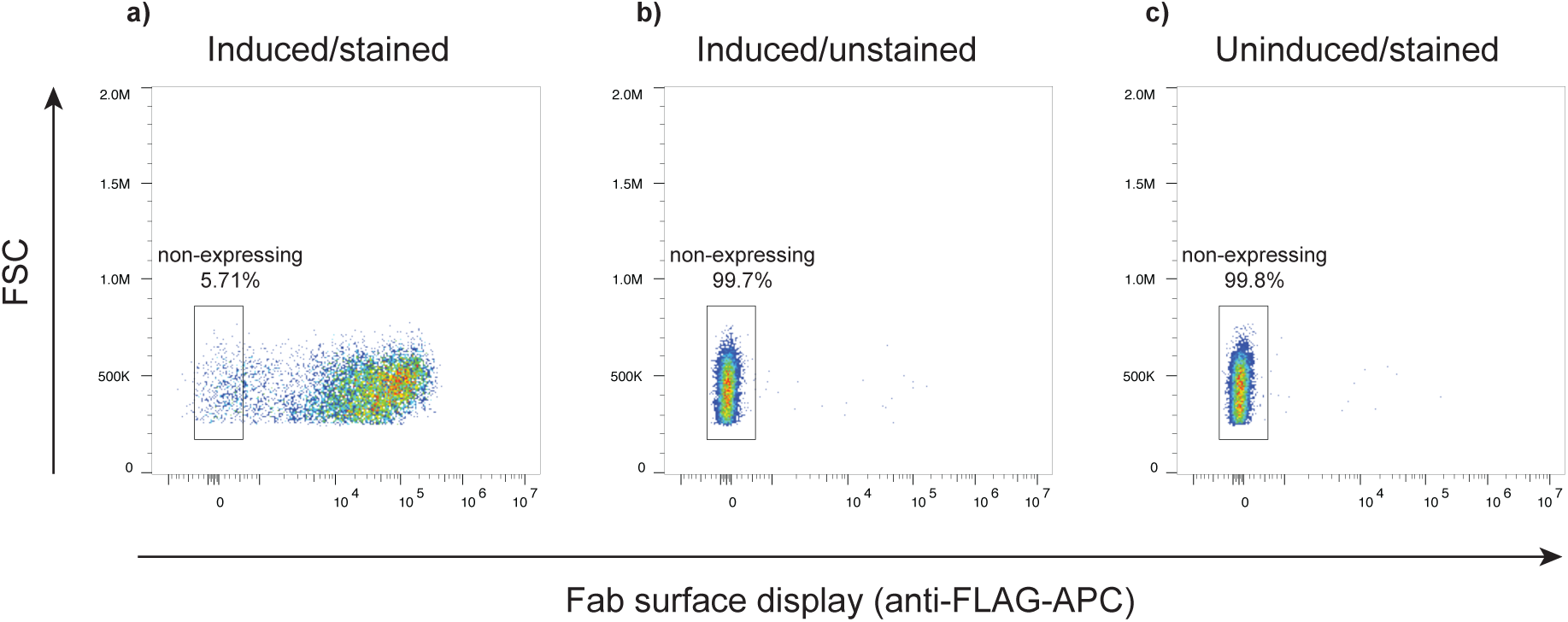
Display of naïve library. a) induced and stained library, b) induced and unstained library and c) uninduced and stained library.

**Supplementary Fig. 8.**
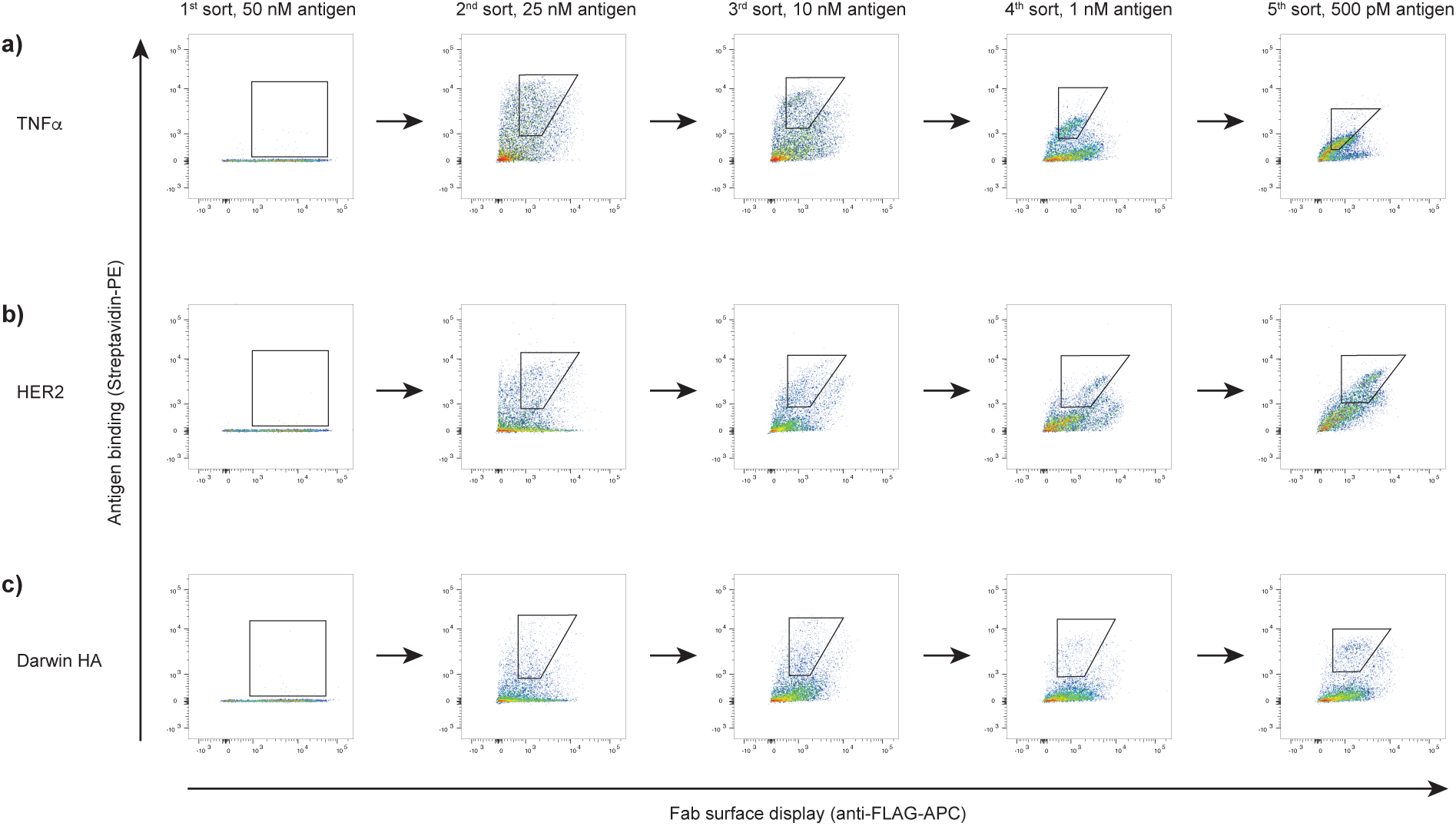
Sort overview. Dot plots for the successive rounds of sorting for a) TNFα, b) HER2 and c) Darwin HA.

**Supplementary Fig. 9.**
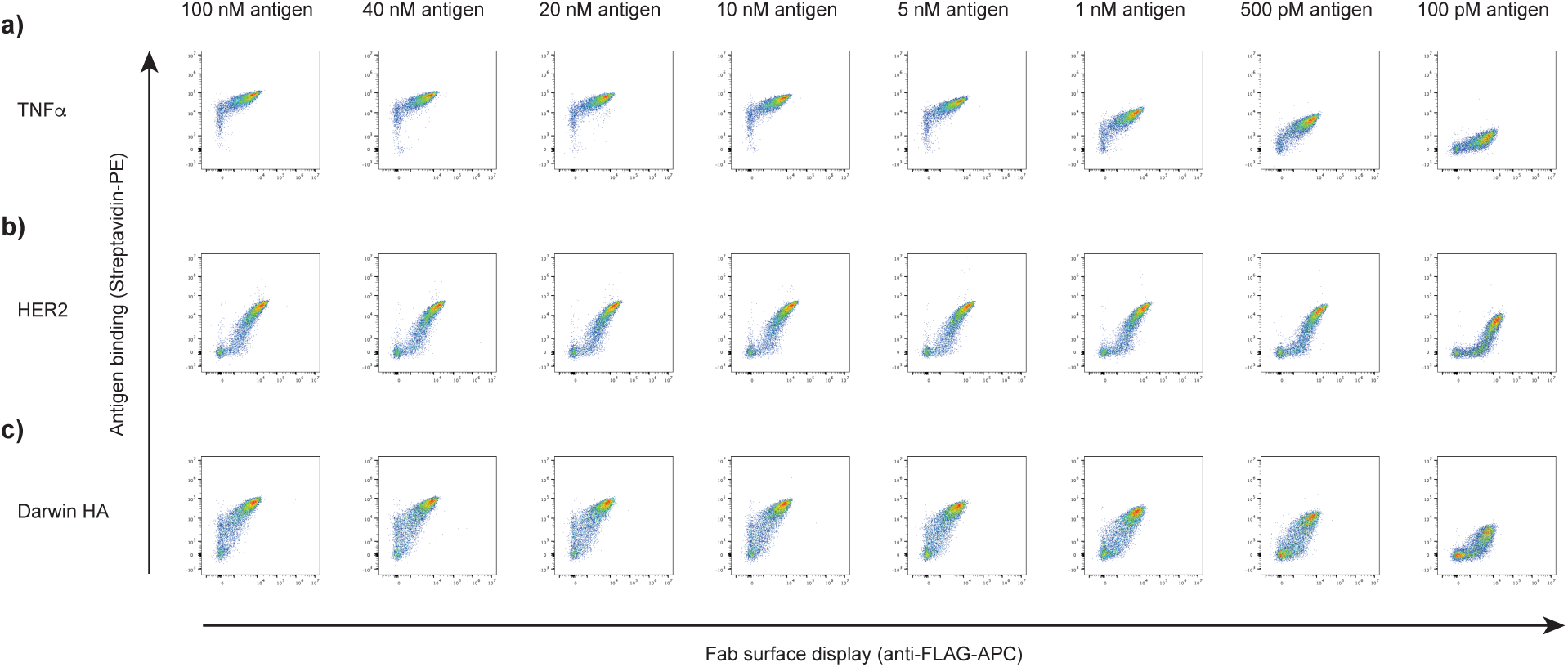
Affinity titration. Antigen titration for a) TNFα mAb, b) HER2 mAb and, c) Darwin HA mAb.

**Supplementary Fig. 10.**
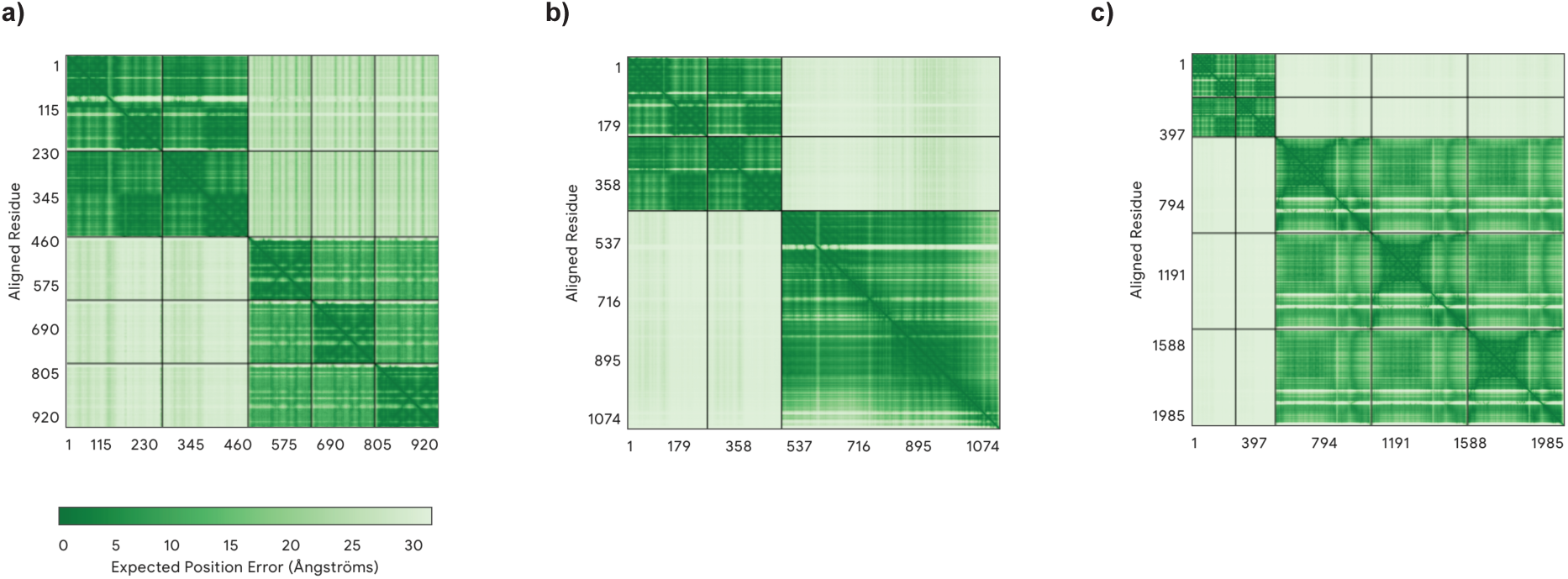
Structural predictions for antibody-antigen interaction. Visualization of the Predicted Aligned Error (PAE) for a) TNFα-mAb, b) HER2-mAb and, c) Darwin HA-mAb.

